# Effect of *Lantana camara* ethanolic leaf extract on survival and migration of MDA-MB-231 triple negative breast cancer cell line

**DOI:** 10.1101/2023.02.04.527114

**Authors:** Arundhaty Pal, Sourav Sanyal, Sayantani Das, Tapas Kumar Sengupta

## Abstract

**Introduction:** Breast cancer is a leading cause of cancer-related death worldwide. *Lantana camara* has been reported to cure a number of ailments, with few studies showing its cytotoxic effects on breast cancer cells. However, the impact of *Lantana camara* on triple negative breast cancer cells is largely obscure to date. The present study investigated the effect of ethanolic extract of *Lantana camara* leaves on the triple negative breast cancer cell line, MDA-MB-231.

**Methods:** Cytotoxic effect of the extract on the cells was determined by cell survival assay. Cell cycle phase distribution was analysed using flow cytometry. To study the effect on nuclear morphology, nuclear stained cells were visualized using epifluorescence microscopy. The induction of apoptosis and generation of reactive oxygen species was conducted using flow cytometry. Cellular migration was studied by performing wound healing assay. Real-time polymerase chain reaction was done to determine the mRNA levels of some key genes.

**Results:** *Lantana camara* leaf extract induced cytomorphological changes and growth inhibitory effect on MDA-MB-231 cells in a dose-dependent manner. The extract also induced G0/G1 cell cycle arrest and nuclear condensation. Flow cytometry analysis confirmed cell death by apoptosis. *Lantana camara* leaf extract also reduced the migration of MDA-MB-231 cells. mRNA expression levels also supported the above observations.

**Conclusions:** This study demonstrates the efficacy of the extract to induce growth-inhibitory and anti-migratory effects on MDA-MB-231 cells. Our results thus suggest that *Lantana camara* leaf extract can be considered as a potent source of chemotherapeutic agents for triple negative breast cancer.

## 1. Introduction

Cancer, as a major cause of death and a significant obstacle to increased life expectancy, accounted for approximately 19.3 million new cases and an estimated 10.0 million deaths worldwide in 2020. With 2.3 million new cases (11.7% of all cancer cases), breast cancer is currently the largest cause of cancer incidence and the fifth leading cause of cancer-related deaths worldwide (Sung et al., 2021).

Breast cancer can be further classified into different molecular subtypes based on immunohistochemical staining: luminal A (ER/PR^+^, HER-2^−^, Ki-67^+^ < 20%), luminal B (ER/PR^+^ < 20%, HER-2^−^, Ki-67^+^ ≥ 20%), HER-2^+^ B2 (ER/PR^+^, HER-2 overexpression), HER-2 overexpression (ER^−^, PR^−^, HER2 overexpression), basal-like Triple Negative Breast Cancer (ER^−^, PR^−^, HER2^−^), and other special subtypes (Goldhirsch et al., 2013). Triple negative breast cancer (TNBC) typically has a higher tumor grade, greater tumor size, higher invasiveness and rate of metastasis, and more frequent recurrence compared to other types of breast cancer (Liang et al., 2021). TNBC is generally common in premenopausal young women under 40, who make up about 15% to 20% of all breast cancer patients (Morris et al., 2007). TNBC patients have a reduced life expectancy, and 40% of them die within the first five years of diagnosis (Dent et al., 2007).

TNBC is resistant to endocrine therapy and molecular targeted therapy because of its unique molecular profile. Surgical removal, radiation therapy, and chemotherapy are currently used in the treatment of TNBC. Bevacizumab and chemotherapy medications have been utilized in combination to treat TNBC in various countries (Yin et al., 2020). However, these treatment options have limitations due to lower efficacy, unwanted side effects, and higher costs. Moreover, a considerable number of cancer patients develop resistance to chemotherapy (Housman et al., 2014) and tumor reappearance is eventually caused by the remaining metastatic lesions. Therefore, a top priority is to find more effective, safe, and affordable treatments against TNBC.

Natural products, mainly plant-derived phytochemicals play a key role in terms of therapeutics. Currently, plant-based anti-cancer drugs account for three-quarters of all prescribed medicines. Taxol, vinblastine, vincristine, podophyllotoxin, and camptothecin are some of the most well-known examples (Soumya et al., 2021). *Curcuma longa*, *Withania somnifera*, *Zingiber officinale*, *Hypericum perforatum*, *Camellia sinensis*, and other medicinal herbs have also shown optimistic anti-breast cancer potential in the research phase (Gospodinova et al., 2021). Despite all of these efforts, clinical usage of plant-based chemotherapeutic medicines has been found to fall short of expectations. Thus, there is an ongoing demand for new, effective, and affordable medications that are free of harmful side effects. A ray of hope is that only 5–15% of the 250,000 higher plants have been screened for therapeutic potential (Soumya et al., 2021). In this scenario, exploiting the vast richness of unfathomed plants in a tropical country like India is both relevant and reasonable. Bioactive compounds derived from these plants can be treatment options for TNBC.

*Lantana camara*, also known as red or wild sage, is an invasive weed of the Verbenaceae family. More than 60 different countries of tropical, subtropical, and temperate regions are home to this flowering species (Ghisalberti, 2000). All the parts of *L. camara*, especially leaves have been utilized in herbal medicines for generations to treat various illnesses such as headache, fever, skin itches, ulcers, rheumatism, measles, chicken pox, asthma, and leprosy (Hernández et al., 2003; Mahdi-Pour et al., 2012). *L. camara* leaves also show anti-bacterial, anti-fungal, nematocidal, insecticidal, and anti-oxidant properties (Kumar and Sivaprakasam, 2007). Altabbaa et al. reported the anti-bacterial activity of Chitosan-coated zinc oxide nanocomposites of *Lantana camara* (Altabbaa et al., 2023). In another study, Amoah et al. found that *Lantana camara* leaf extract ameliorates neuroinflammation and memory deficit associated with scopolamine-induced Alzheimer’s-like cognitive impairment in zebrafish and mice (Amoah et al., 2023). Different types of terpenes such as germacrene-D, alpha-copaene, b-curcumene, lantadenes, lantanilic acid, oleanolic acid, ursolic acid, etc are some notable constituents of the *Lantana camara* leaves (Kazmi et al., 2012). Various glycosides, proteins, carbohydrates, and fatty acids are also found in *L. camara* leaves (Kazmi et al., 2012). Wu and his coworkers further characterized various other euphane-type triterpenoids from methanolic extract of aerial parts of *L. camara* which showed anti-inflammatory effects ((Wu et al., 2020). Various studies have also reported that different extracts of *L. camara* leaves and their bioactive compounds display anti-cancer activity. Human lung carcinoma cell line (A549) and mouse melanoma (B16F10) revealed *in vitro* cytotoxicity when exposed to *L. camara* leaf methanolic extract (Hariharapura et al., 2004). Another study has reported that ethanolic extract of *L. camara* inhibited proliferation and induced apoptosis of MCF-7 breast cancer cell line (Han et al., 2015). Shamsee et al. identified four pentacyclic triterpenoids (lantadene-A, lantadene-B, lantadene-C, and icterogenin) from the methanolic extract of *L. camara* leaves and found that all these compounds can reduce MCF-7 cell viability in a dose-dependent manner (Shamsee et al., 2019). Lantadene-B showed the most potent anti-cancer activity among these pentacyclic triterpenoids and induced G0/G1 cell cycle arrest in MCF-7 cells (Shamsee et al., 2019). In another study, Jaafar et al. reported that Lantadene A-loaded gold nanoparticles could induce genotoxicity and cytotoxicity against MCF-7 cells (Jaafar et al., 2020). Similarly, Hublikar et al. found that silver nanoparticles synthesized using *Lantana camara* leaf extract have cytotoxic activity on A549 and MCF-7 cell lines (Hublikar et al., 2023). Although the effect of *L. camara* extracts was studied on different cancer cell lines, only a few studies are there so far that investigated the effect of *L. camara* extract on triple negative breast cancer cells. Ramkumar et al. reported cytotoxic properties of *L. camara* root extract-derived gold nanoparticles on MDA-MB-231 cells (Ramkumar et al., 2017) and in a recent study, El-Din et al. showed that methanolic extract of *L. camara* leaves could exert cytotoxic effect on several cancer cell lines including MCF-7 and MDA-MB-231 breast cancer cells; however, the mechanism of cytotoxicity in breast cancer cell lines was not investigated (El-Din et al., 2022). More recently, Mishra et al reported that extracts of the whole *Lantana camara* plant harbours cytotoxic effect against triple negative MDA-MB-231 cells although the mechanism of cytotoxicity was unexplored for this cell line (Mishra et al., 2023). It is therefore important to explore the mode of cytotoxic action of *Lantana camara* leaf extract on breast cancer cell lines, particularly on the triple negative breast cancer cells. It is also noteworthy that the level and kind of phytochemicals present in various *Lantana* species are reported to be quite varied (Sharma et al., 1991). Therefore, in the present study, we investigated the effects of *L. camara* (from Mohanpur, Nadia, West Bengal, India) leaf ethanolic extract on two important hallmarks of cancer, cell death and migration in the triple negative breast cancer cell line MDA-MB-231.

## 2. Materials and methods

### 2.1 Cell culture

MDA-MB-231 and MCF-7 cell lines were obtained from National Centre for Cell Science (NCCS), Pune, India. HEK293T (human kidney epithelial-like) cells were kindly gifted by Dr. Sumit Sen Santara (DBS, IISER Kolkata). All cell lines were maintained in Dulbecco’s Modified Eagle Medium (DMEM) High Glucose (HiMedia, India) supplemented with 10% (v/v) Fetal Bovine Serum (FBS) (HiMedia, India), 10000 U/mL Penicillin and 10 mg/mL Streptomycin antibiotic solution (HiMedia, India). Cells were incubated at 37°C in 95% air in a humidified incubator with 5% CO_2_. 1X Trypsin-EDTA solution (HiMedia, India) was used for cell harvesting during sub-culturing.

### 2.2 *Lantana camara* leaf extract preparation

Healthy leaves of *Lantana camara* were collected from the campus and surroundings of the Indian Institute of Science Education and Research Kolkata (latitude-22°57´50´´N and longitude-88°31´28´´E). The leaves were identified and authenticated by an expert botanist of Central National Herbarium, Botanical Survey of India, West Bengal, India. A voucher specimen was deposited at their herbarium with a voucher/reference number IISER/AP/01. The collected plant leaves were rinsed with tap water, followed by distilled water, and then shade-dried at 30°C for 48 hours. Dried leaves were ground into a fine powder using mortar and pestle, and then subjected to extraction with absolute ethanol in a ratio of 1:10 (w/v) by gentle mixing on an orbital shaker at 37°C for 18 hours. The extracted mixture was filtered using Whatman No.2 filter paper and the filtrate was concentrated using a rotary evaporator under low pressure and 37°C temperature. The resultant extract residues were then collected and dissolved in molecular-grade ethanol (Merck, Germany) to prepare stock solution as required. This stock solution was filtered using a syringe filter (0.22 microns) and was kept at 4°C for future use.

As *Lantana camara* leaves were extracted with ethanol and the cells were treated with the ethanolic extracts, first the effect of different concentrations (percentage) of ethanol on MDA-MB-231 cell lines was determined in order to select a fixed final concentration of ethanol with a minimal effect on the cells as the vehicle in all treatments. It was also important to determine the ethanol concentration dependent solubility of the extract. Considering both the concerns, solvent tolerance assay was performed and it was found that 0.8% (v/v) ethanol as the final concentration in all experimental treatment as most appropriate (Supplementary Figure S1). In order to treat the cells with different amounts/doses of the extract but keeping the final concentration of ethanol as the solvent/vehicle as 0.8%, we had diluted the stock solution (50 mg/mL) of the extract in the molecular grade ethanol to make working stocks for respective treatments and a fixed volume of the respective working stocks was added to cell culture media for all treatments.

### 2.3 Cell morphology analysis

MDA-MB-231 cells were seeded in 35 mm dishes (0.15 x10^6^ cells/dish) and incubated overnight. The spent medium was then discarded and cells were treated with 40 µg/mL, 80 µg/mL, 120 µg/mL and 150 µg/mL of the extract for 24 hours. Cells treated without any extract or ethanol and cells treated with 0.8% ethanol were kept as untreated control and vehicle control, respectively. Cell morphology was examined and images were captured using an inverted microscope (IX81, Olympus, Japan) with a camera (Photometrics, CoolSNAP MYO) under 40X magnification.

### 2.4 Cell viability assay

The cytotoxicity of *L. camara* was tested on MDA-MB-231, HEK293T and MCF-7 cells by 3-(4,5-dimethylthiazol-2-yl)-2,5-diphenyltetrazolium bromide (MTT) colorimetric assay. Cells were seeded in a 96-well plate (0.5x10^4^ cells/well) and incubated overnight. Cells were then treated with 20 µg/mL, 40 µg/mL, 60 µg/mL, 80 µg/mL, 100 µg/mL, 120 µg/mL, 140 µg/mL, 160 µg/mL and 180 µg/mL of the extract for 24 hours. Cells treated with 0.8% ethanol were kept as vehicle control. After treatment, spent media from the wells was discarded, fresh complete media with a final concentration of 0.5 mg/mL of MTT solution (SRL, India) was added to each well and the cells were incubated in the dark at 37°C and 5% CO_2_ for 4 hours. Thereafter, the MTT containing medium was carefully aspirated from the plate, 100 µL of dimethyl sulfoxide (DMSO) was added to each well and incubated for 30 minutes in the dark at 37°C. Then absorbance was measured in a microplate reader (Biotek, Agilent technologies, CA, USA) using Gene5 software at 595 nm. The cytotoxic effect of the leaf extract was expressed as viability percentage in comparison to vehicle control. Dose response curve was created by using GraphPad Prism 6.01 (GraphPad software, CA, USA).

### 2.5 Cell cycle phase distribution assay

MDA-MB-231 cells were seeded in 35 mm cell culture dishes (0.15 x10^6^ cells/dish) and incubated overnight. Cells were then treated with 60 μg/mL, 80 μg/mL, 100 μg/mL, 120 μg/mL doses of the leaf extract along with vehicle treated and untreated control cells. Following 24 hours of treatment, cells were harvested from the culture dishes by treating with 1X Trypsin-EDTA and collected along with the floating cells derived from spent media. Cell pellets were washed with ice-cold 1X phosphate buffered saline (PBS). Then the cells were fixed with 70% ethanol and incubated at 4°C overnight. This was followed by washing the cells with ice-cold 1X PBS and resuspension in 500 µL propidium iodide (PI) solution [475 µL 1X PBS+20 µL PI (HiMedia, India) from 1 mg/mL stock+5 µL RNase A (HiMedia, India) from 20 mg/mL stock)] and incubated at 37°C for 20 minutes in dark. Thereafter, the PI solution was removed and cells were resuspended in 500 µL 1X PBS and the samples were analysed using BD FACS Fortessa flow cytometer (BD Biosciences, USA). Cell cycle distribution was (Sub-G1, G1/G0, S and G2/M) determined and the graph was plotted by using GraphPad Prism 6.01 (GraphPad software, CA, USA).

### 2.6 Nuclear morphological analysis by 4’,6-diamidino-2-phenylindole (DAPI) staining

MDA-MB-231 cells (0.15 x10^6^ cells/well) were seeded in a 24-well plate containing coverslips and then treated with 60 μg/mL, 120 μg/mL, 150 μg/mL and 180 μg/mL doses of the leaf extract along with vehicle treated and untreated control cells for 24 hours. Following treatment, media was removed and the wells (with coverslips) were washed with 1X D-PBS (1X PBS with 0.9mM calcium chloride and 0.5 mM magnesium chloride). Cells were further fixed by adding ice-cold 4% paraformaldehyde and incubated for 15 minutes at room temperature. After fixation, cells on the coverslips were washed with 1X PBS and then permeabilized by adding permeabilization buffer [1X D-PBS+0.5% Tween 20] and incubated for 20 minutes. Then the cells were washed with 1X PBS, 1 µg/mL of DAPI solution (in 1X PBS) was added to each well (with coverslip) and incubated for 15 minutes in the dark. Cells were again washed with 1X PBS; coverslips were removed from the wells, mounted on clean glass slides using 5% glycerol as mounting agent and sealed. The slides were observed and images were captured under an inverted microscope (IX81, Olympus, Japan) with a camera (Photometrics, CoolSNAP MYO).

### 2.7 Annexin V-FITC/PI double staining

Apoptotic and non-apoptotic cells were differentiated using the Annexin V-FITC Apoptosis detection kit (BD Biosciences, USA) according to the manufacturer’s protocol. MDA-MB-231 cells (0.2 x10^6^ cells/dish) were seeded in 35 mm culture dishes and then 60 μg/mL, 120 μg/mL, 180 μg/mL doses of the leaf extract along with vehicle treated cells for 24 hours. Following treatment, cells were harvested from the culture dishes by treating with 1X Trypsin-EDTA and collected along with the floating cells derived from spent media. Cell pellet was once washed with ice-cold 1X PBS and then resuspended in 100 μL of 1X binding buffer. 5 μL each of Annexin V-FITC and Propidium iodide was added for staining and gently vortexed followed by incubation in the dark for 15 minutes at room temperature. Finally, 400 μL of 1X binding buffer was added to each tube and the samples were analysed by using BD FACS Fortessa flow cytometer (BD Biosciences, USA). The percentage of apoptotic and non-apoptotic cell population was determined by evaluating the relative amount of Annexin V-FITC positive alone cells (early apoptosis), both Annexin V-FITC and PI-positive cells (late apoptosis) and PI positive alone cells (non-apoptotic). Cell population negative for both Annexin V-FITC and PI was considered as viable cells. Data was represented as scatter plots displaying the distribution of cells after Annexin V-FITC/ PI double staining, where lower left quadrant represents viable cells, lower right quadrant represents early apoptotic population, late apoptotic population is reflected in upper right and non-apoptotic population is represented in upper left quadrant. Data was also represented graphically by using GraphPad Prism 6.01 (GraphPad software, CA, USA).

### 2.8 Wound healing assay

MDA-MB-231 cells (0.2 x10^6^ cells/dish) were seeded in 35 mm dishes and grown until a uniform monolayer of cells was formed. Once a monolayer was formed, a wound was created in each dish using a sterile P10 microtip and the spent media was removed from the culture dish. The monolayer of cells was washed twice with 1X PBS, complete media was added to the cells followed by addition of cycloheximide (0.5 μg/mL). Immediately after addition of cycloheximide, the wounded cells were treated with 40 μg/mL, 60 μg/mL and 80 μg/mL doses of the leaf extract along with vehicle treated and untreated control cells. The cells were imaged at 0 hour post treatment using inverted microscope (IX81, Olympus, Japan) with a camera (Photometrics, CoolSNAP MYO). The cells were incubated at 37°C in 95% air in a humidified incubator with 5% CO_2_ and images were intermittently captured at 12 hours and 24 hours post treatment. The images were analysed and wound closure was calculated by ImageJ 1.54b (ImageJ software, NIH, USA) using the following formula:

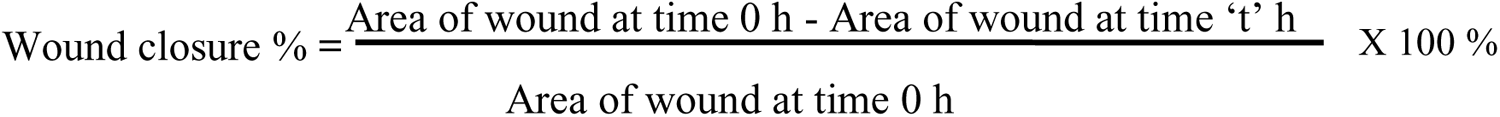

where t = specific time intervals after 0 hour

Data was represented graphically by using GraphPad Prism 6.01 (GraphPad software, CA, USA).

### 2.9 RNA isolation and reverse transcription quantitative real-time polymerase chain reaction (RT-qPCR)

MDA-MB-231 cells were cultured in 60 mm dishes for up to 70-80% confluency. Then the cells were treated with either vehicle (0.8% ethanol) or 40 µg/mL, 60 µg/mL, 120 µg/mL and 180 µg/mL doses of *Lantana camara* leaf ethanolic extract for 24 hours. Total RNA was isolated from vehicle and *Lantana camara* leaf ethanolic extract-treated MDA-MB-231 cells using RNAiso Plus (Takara, Japan) as per the manufacturer’s protocol. The RNA samples were treated with RNase-free DNase I (Thermo Fisher Scientific, USA) to digest any residual chromosomal DNA and then quantified using a microplate spectrophotometer (BioTek Epoch Microplate Spectrophotometer, Agilent Technologies Inc, USA).

These RNA samples were used for cDNA synthesis using PrimeScript 1st strand cDNA synthesis kit (Takara, Japan) as per the manufacturer’s instructions. Briefly, a total of 10 µL of reaction volume containing 1 µg of RNA, 2 µL of 50 µM Random 6-mers, and 1 µL of 10 mM dNTP mixture were incubated at 65°C for 5 minutes and immediately kept on ice. 4 µL of 5X PrimeScript buffer, 0.5 µL of 40 U/µL RNase inhibitor and 1 µL of 200 U/µL PrimeScript reverse transcriptase were added to the initial 10 µL reaction volume, and the final volume was made up to 20 µL with sterile nuclease-free water. The reaction mixtures were first incubated at 30°C for 10 minutes, then at 42°C for 60 minutes, and finally at 70°C for 15 minutes using Veriti 96 well thermal cycler (Applied Biosystems, Thermo Fisher Scientific, USA). The resulting products were kept at -20°C until further use.

Real-time PCR was carried out using a TB Green Premix Ex Taq (Tli RNaseH Plus) kit (Takara, Japan) according to the manufacturer’s protocol. Briefly, 25 ng of cDNA, along with 5 µL of 2X TB Green Premix Ex Taq (RNaseH Plus) and 0.2 µL of each 10 µM forward and reverse primer were added to a total reaction volume of 10 µL, and subjected to thermo-cycling in a CFX96 Touch Real-time PCR Detection System (Bio-Rad Laboratories, Hercules, CA, USA). The conditions for thermal cycling were as follows: 1 cycle of initial denaturation at 95°C for 30 seconds, then 40 cycles of 95°C for 5 seconds, 55°C for 30 seconds (annealing), and 72°C for 30 seconds (extension). The dissociation curves were built right after the PCR run to check and justify the results. Primers of Cyclin D1, p21, Bcl-2, Bax, N-cadherin, and Vimentin were used for gene expression study. All the primers were designed for this study (Supplementary Table S1) and checked by the Primer-BLAST tool of the National Institute of Health (NIH, Bethesda, Maryland, USA). Gene expression was normalized to 18S rRNA housekeeping gene. Data were analysed using the 2^-ΔΔCt^ method and plotted as fold change versus vehicle control (Livak and Schmittgen, 2001).

### 2.10 Statistical analysis

All experiments were conducted for at least three times and data was represented as mean ± SEM (Standard Error of Mean). Data was plotted using GraphPad Prism 6.01 (GraphPad software, CA, USA) and statistical analysis was done by performing parametric Student’s t-test. *P*<0.05 was considered to indicate a statistically significant difference.

## 3. Results

### 3.1 *L. camara* leaf ethanolic extract induced cytotoxic effects in MDA-MB-231 cells

MDA-MB-231 cells have spindle-shaped morphology under normal culture conditions. To evaluate whether the leaf extract induced any morphological changes, MDA-MB-231 cells were exposed to 40 μg/mL, 80 μg/mL, 120 μg/mL and 150 μg/mL of leaf extract, and vehicle (0.8% ethanol) for 24 hours and the observed changes in the morphology of the cells are presented in Figure 1A. It was observed that at lower doses (40 μg/mL and 80 μg/mL) of the extract, MDA-MB-231 cells were shorter in length and cell shrinkage was observed at 80 μg/mL dose as compared to the control conditions (vehicle treated and untreated cells). Further, it was observed that at the higher doses (120 μg/mL and 150 μg/mL), MDA-MB-231 cells appeared more spherical and the majority of such spherical cells were found to lose their adherent property, thus became floating in the growth media.

**Figure 1.**
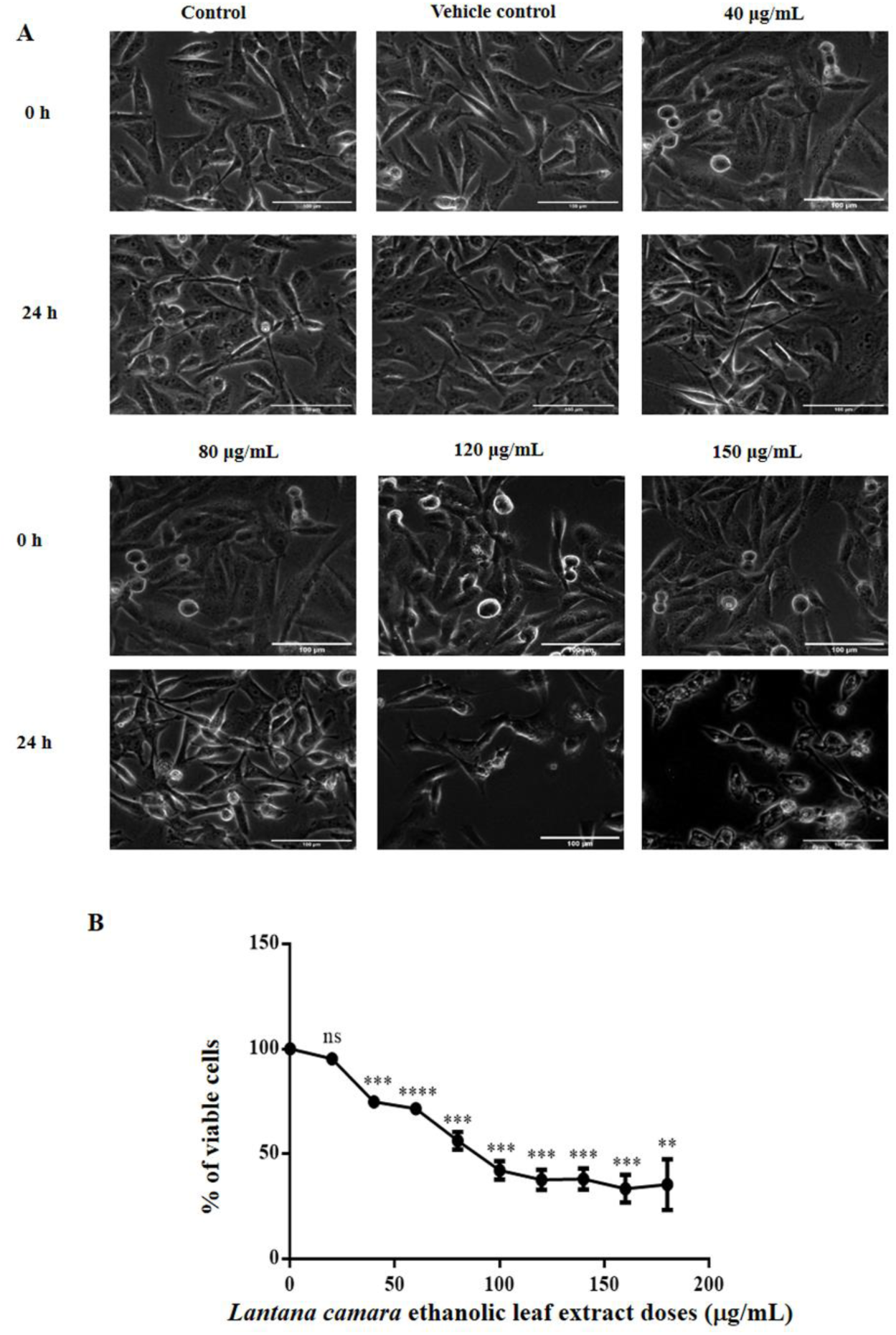
*L. camara* leaf ethanolic extract produced cytotoxic effects in MDA-MB-231 cells. **(A)** MDA-MB-231 cells treated with different concentrations of the extract for 24 hours, and images were captured at 0 hours and 24 hours after treatment. **(B)** MDA-MB-231 cells were treated with different concentrations of the extract for 24 hours, and the viability of the cells was evaluated by MTT assay. Scatter plot distribution is represented as means ± SEM of three independent biological replicates where * is (*P*≤0.05), ** (*P*≤0.01), *** (*P*≤0.001) and **** (*P*≤0.0001) and ns signifies non-significant.

The observed induction of morphological changes was indicative of cytotoxic effect being exerted by the extract on the MDA-MB-231 cells. To understand whether altered morphology is due to the induction of cytotoxicity by leaf extract, MDA-MB-231 cells were treated with various doses (20 μg/mL to 180 μg/mL) of the leaf extract for 24 hours and the viability of the treated MDA-MB-231 cells was evaluated using MTT assay. The graph in Figure 1B represents the dose-dependent viability of MDA-MB-231 cells where the viability of vehicle-treated cells was considered to be 100%. It was observed that leaf extract impeded the viability of the cells in a dose-dependent manner and the half-maximal inhibitory concentration (IC_50_) of the extract was found to be 111.33 μg/mL.

The specificity of *Lantana camara* leaf extract towards the MDA-MB-231 cells were tested by comparing the cytotoxic activity of the extract towards the normal kidney epithelial-like HEK293T cells (Supplementary Figure S2) and its was observed that the extract was less toxic towards the normal cells.

### 3.2 Induction of G0/G1 cell cycle arrest by *L. camara* leaf extract in MDA-MB-231 cells

To further evaluate the mechanism of the anti-proliferative effect of the extract on MDA-MB-231 cells, cell-cycle phase distribution was analysed by using flow cytometry (Figure 2). Figure 2A is a representative image of the distribution of MDA-MB-231 cells in the different phases of cell cycle (Sub-G1, G0/G1, S and G2/M) when treated for 24 hours with 60 μg/mL, 80 μg/mL, 100 μg/mL and 120 μg/mL of the extract along with vehicle treated and untreated control. This range of doses of *Lantana camara* extract for the cell cycle analysis was selected based on the results of MTT assay where significant reduction in cell viability was observed for 24 hours of treatment and also by considering the fact that cell cycle arrest is an early event compare to cell death. Analysis of the results (Figure 2B) revealed that G0/G1 population of MDA-MB-231 cells increased from 61.35 ± 2.6 % (vehicle control) to 78.80 ± 1.58 %, 79.68 ± 2.07%, 78.15 ± 2.17 % and 75.98 ± 3% when treated with 60 μg/mL, 80 μg/mL, 100 μg/mL and 120 μg/mL of leaf extract, respectively. This finding implied that treatment with *L. camara* leaf extract induced arrest of MDA-MB-231 cells at the G0/G1 phase and the effect remained essentially unaltered with increase in dose. The observed increase in the G0/G1 population was also accompanied by significant concomitant decrease in S-phase and G2/M phase population at all doses in *L. camara* extract-treated cells as compared to control conditions. In the sub-G1 population, slight increase in the percentage of cell population, due to the treatment with leaf extract was also observed, although the differences were statistically insignificant. Taken together, our result established that *L. camara* leaf extract induced dose-independent G0/G1 arrest in MDA-MB-231 cells accompanied by reduction in S and G2/M population as compared to vehicle control cells.

**Figure 2.**
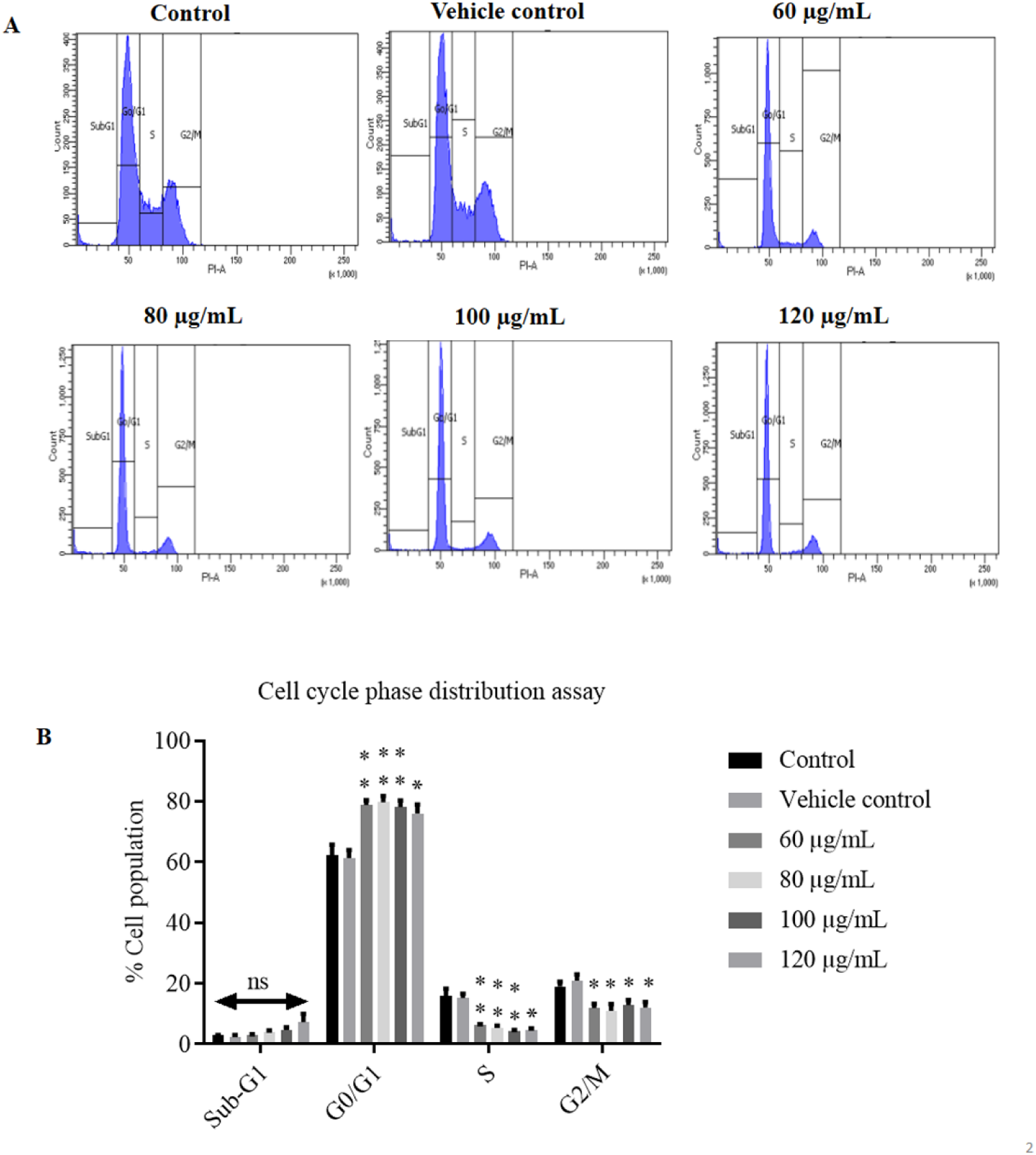
*L. camara* leaf ethanolic extract induced G0/G1 cell cycle arrest in MDA-MB-231 cells. **(A)** Representative images of cell cycle phase distribution of MDA-MB-231 cells following 24 hours of treatment with the leaf extract at different doses. **(B)** Bar diagram showing the percentage of MDA-MB-231 cells in different cell phases (G0/G1, S, G2/M, and sub-G1) of the cell cycle following 24 hours treatment of the extract at different doses. Data are means of three independent experiments and presented as mean ± SEM where * is (*P*≤0.05), ** (*P*≤0.01), *** (*P*≤0.001) and **** (*P*≤0.0001) and ns signifies non-significant.

### 3.3 Nuclear condensation was induced by *L. camara* leaf ethanolic extract

To further understand the underlying mechanism of cytotoxicity imposed by *L. camara*, we analysed chromatin condensation and nuclear fragmentation of MDA-MB-231 cells by staining with the DNA-binding fluorescent dye, DAPI (4’,6-diamidino-2-phenylindole) as nuclear condensation reflects early stages of cell death. Cells were treated with 60 μg/mL, 120 μg/mL, 150 μg/mL and 180 μg/mL of the extract along with vehicle treated and untreated control cells. Increase in sub-G1 population is generally indicative of nuclear fragmentation and cell death, however in experiment 3.2, no significant increase in the sub-G1 population was observed till 120 μg/mL of dose following 24 hours of treatment. Therefore, besides 60 μg/mL and120 μg/mL, two other higher doses (150 μg/mL and 180 μg/mL) were introduced in experiment 3.3 keeping the treatment time (24 hours) same as previous experiment 3.2 for studying nuclear condensation and fragmentation. It was observed (Figure 3) that with increase in the dose of the leaf extract, nuclear condensation and fragmentation was induced in the cells. The corresponding DIC images also corroborated the overall morphological changes induced (as observed earlier in Figure 1A) by treatment with the leaf extract. In contrast, untreated and vehicle treated cells displayed uniform chromatin staining and unaltered cellular morphology.

**Figure 3.**
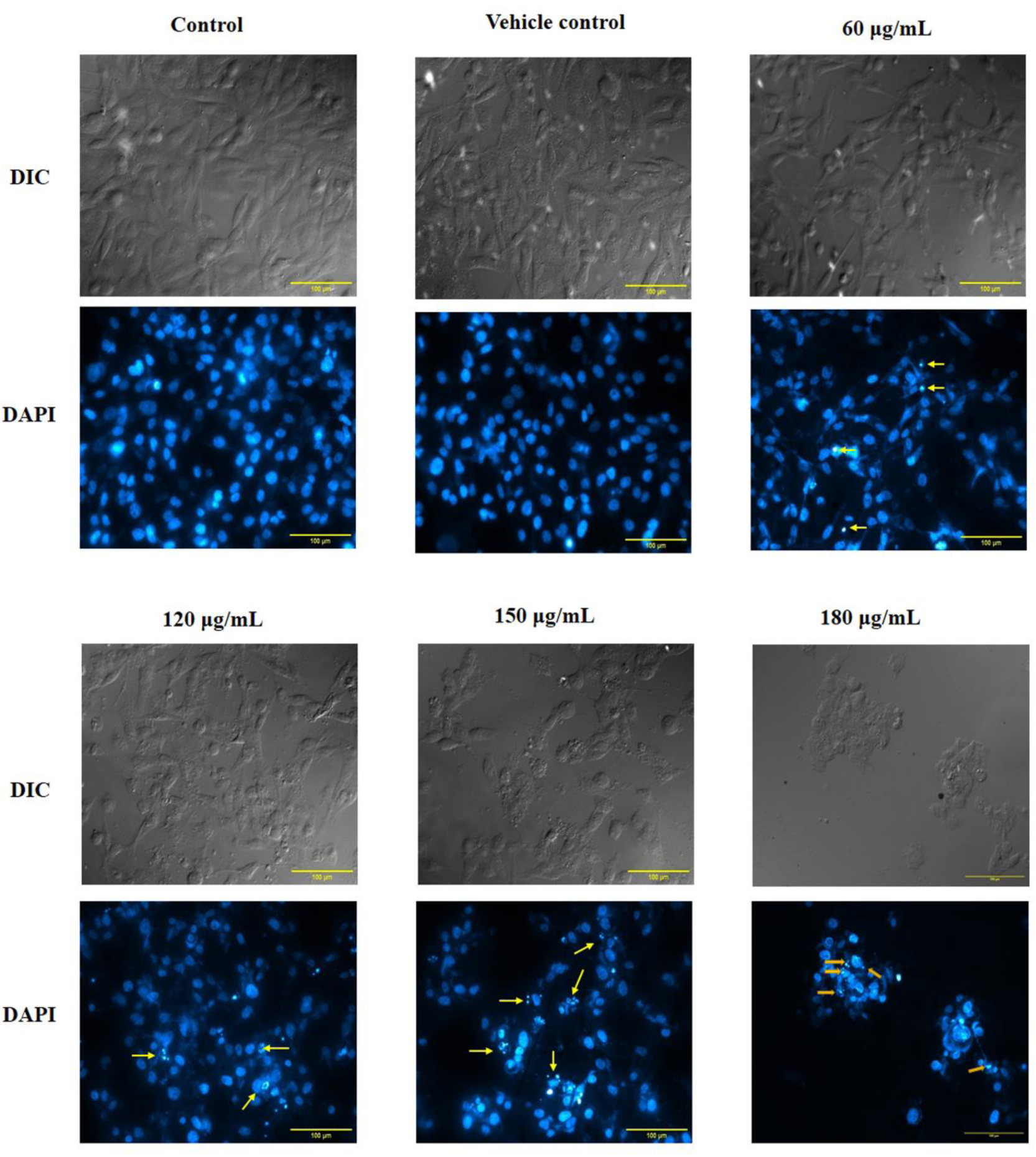
*L. camara* leaf ethanolic extract produced chromatin condensation. MDA-MB-231 cells were treated using different doses of the extract for 24 hours and nuclear staining was performed using DAPI and images were taken. Condensed and fragmented nuclei are marked by arrows. Upper panel: DIC images of MDA-MB-231 for corresponding doses; Lower panel: DAPI images of MDA-MB-231 for corresponding doses.

### 3.4 *L. camara* leaf ethanolic extract promoted apoptosis in MDA-MB-231 cells

The induction of nuclear condensation and fragmentation by the extract was partially indicative of programmed cell death. To elucidate whether the growth inhibitory effect of the extract was due to apoptosis-mediated cell death, MDA-MB-231 cells were exposed to three different doses (60 μg/mL, 120 μg/mL and 180 μg/mL) of the extract selected from the same range of doses used in experiment 3.3 for 24 hours along with vehicle treated cells. Vehicle and extract treated cells were analysed by flow cytometry after double staining with Annexin V-FITC/ PI. Graphical representation in Figure 4B showed that *L. camara* leaf extract induced apoptotic cell death in a dose-dependent manner. Compared to vehicle control, 60 μg/mL of extract-treated cells had a very slight increase in early apoptotic population, late apoptotic and non-apoptotic population. There was a sharp increase in early, late and non-apoptotic populations when the cells were exposed to near IC_50_ dose, i.e., 120 μg/mL. However, the maximum decrease in cellular viability was observed when the cells were treated with 180 μg/mL dose of the extract. The percentage of viable cells decreased from 92.14% ± 0.94% in vehicle-treated cells to 64.67% ± 1.97% in 180 μg/mL extract treated cells. There was a steep increase in early and late apoptotic populations when the cells were treated with 180 μg/mL of the extract. The above findings culminated in the fact that *L. camara* leaf extract induced death in MDA-MB-231 cells mainly by the process of apoptosis.

**Figure 4.**
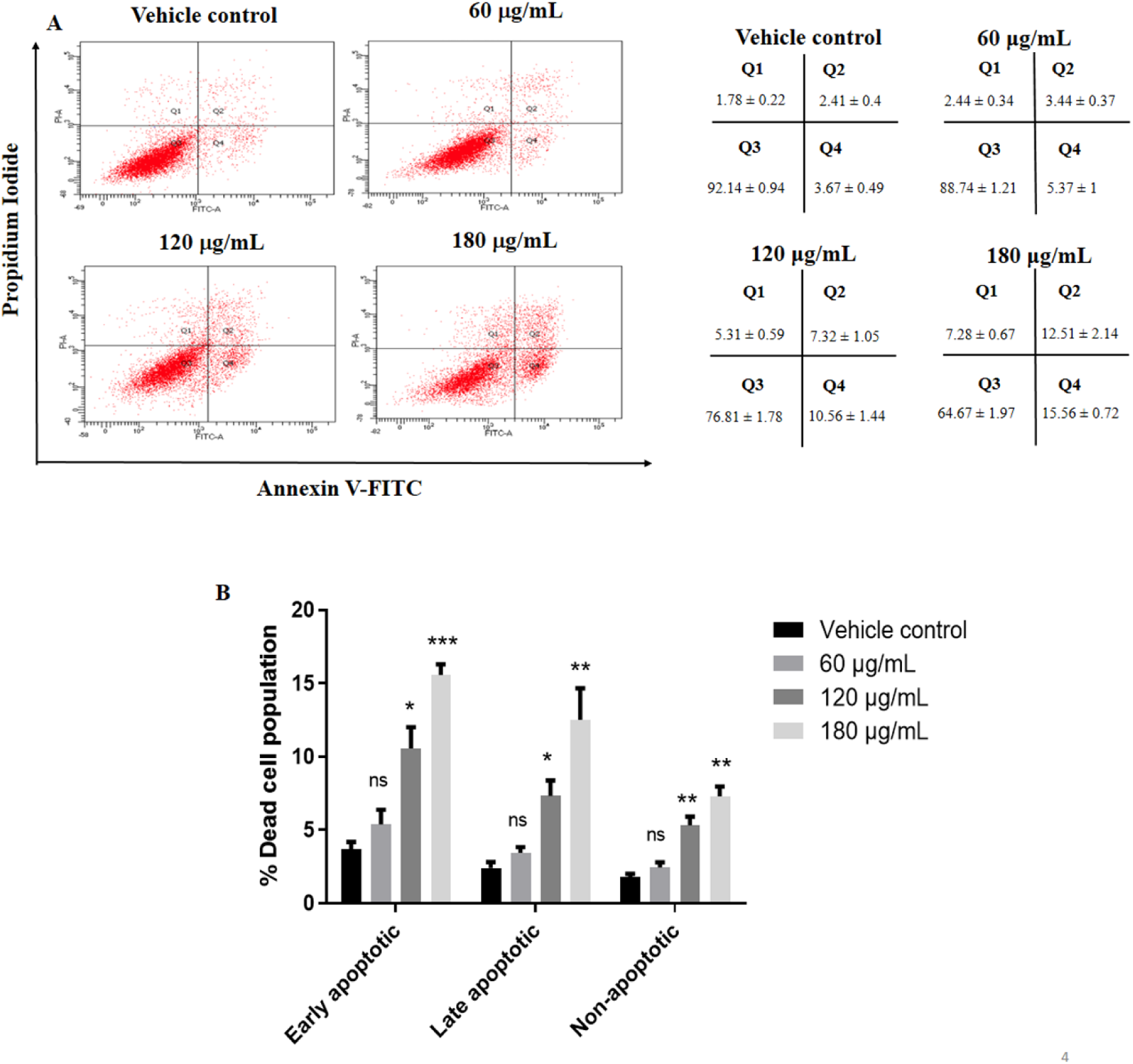
*L. camara* leaf ethanolic extract promoted apoptosis in MDA-MB-231 cells. MDA-MB-231 cells were treated using different doses of the extract for 24 hours followed by double staining the cells with Annexin V-FITC/PI and further analyzed using flow cytometer. **(A)** Left Panel: Scatter analysis with Annexin V-FITC/PI double staining represents the percentage of viable MDA-MB-231 cells (lower left quadrant, Q3) and cells undergoing early apoptosis (lower right quadrant,Q4), late apoptosis (upper right quadrant, Q2) and non-apoptotic (upper left quadrant, Q1) mode of cell death upon treatment with various doses of the leaf extract for 24 hours; Right Panel: Data represents mean ± SEM. of percentage of cells distributed in Q1, Q2, Q3 and Q4. **(B)** Bar diagram represents the percentage of non-viable cell population in early apoptotic, late apoptotic and non-apoptotic phases. Data is represented as mean ± SEM of three independent biological replicates where * is (*P*≤0.05), ** (*P*≤0.01), *** (*P*≤0.001) and ns signifies non-significant.

### 3.5 Migration of MDA-MB-231 cells was impeded by *L. camara* leaf ethanolic extract

The ability of cancer cells to migrate is indispensable for the initiation of metastasis. An efficient therapeutic approach includes restriction of the migratory potential of the cancer cells. To assess the impact of the extract on the migratory potential of MDA-MB-231, cells were treated with lower doses (40 μg/mL, 60 μg/mL and 80 μg/mL) of the extract as at the higher doses, cells lose their morphology and adherent property upon 24 hours treatment, thereby difficult to monitor the cell migration. The wound was given in the presence of 0.5 μg/mL of Cycloheximide. As shown in Figure 5A, in comparison to vehicle control, a lesser wound area was healed by the treated cells. The graphical representation of the wound closure percentage (Figure 5B) indicated that although the wound closure activity was not suppressed significantly after 12 hours of treatment, significant impediment to wound closure activity was observed post 24 hours of treatment. It was seen that the migratory potential of MDA-MB-231 cells was drastically reduced in a dose-independent manner, where the wound closure percentage was 60.27% ± 5.12% for vehicle control, it was found to be 42.08% ± 1.68%, 40.05% ± 3.11% and 38.04% ± 10.38% for 40 μg/mL, 60 μg/mL and 80 μg/mL respectively. Thus, our result demonstrated that *L. camara* leaf ethanolic extract impeded the migratory ability of MDA-MB-231 cells.

**Figure 5.**
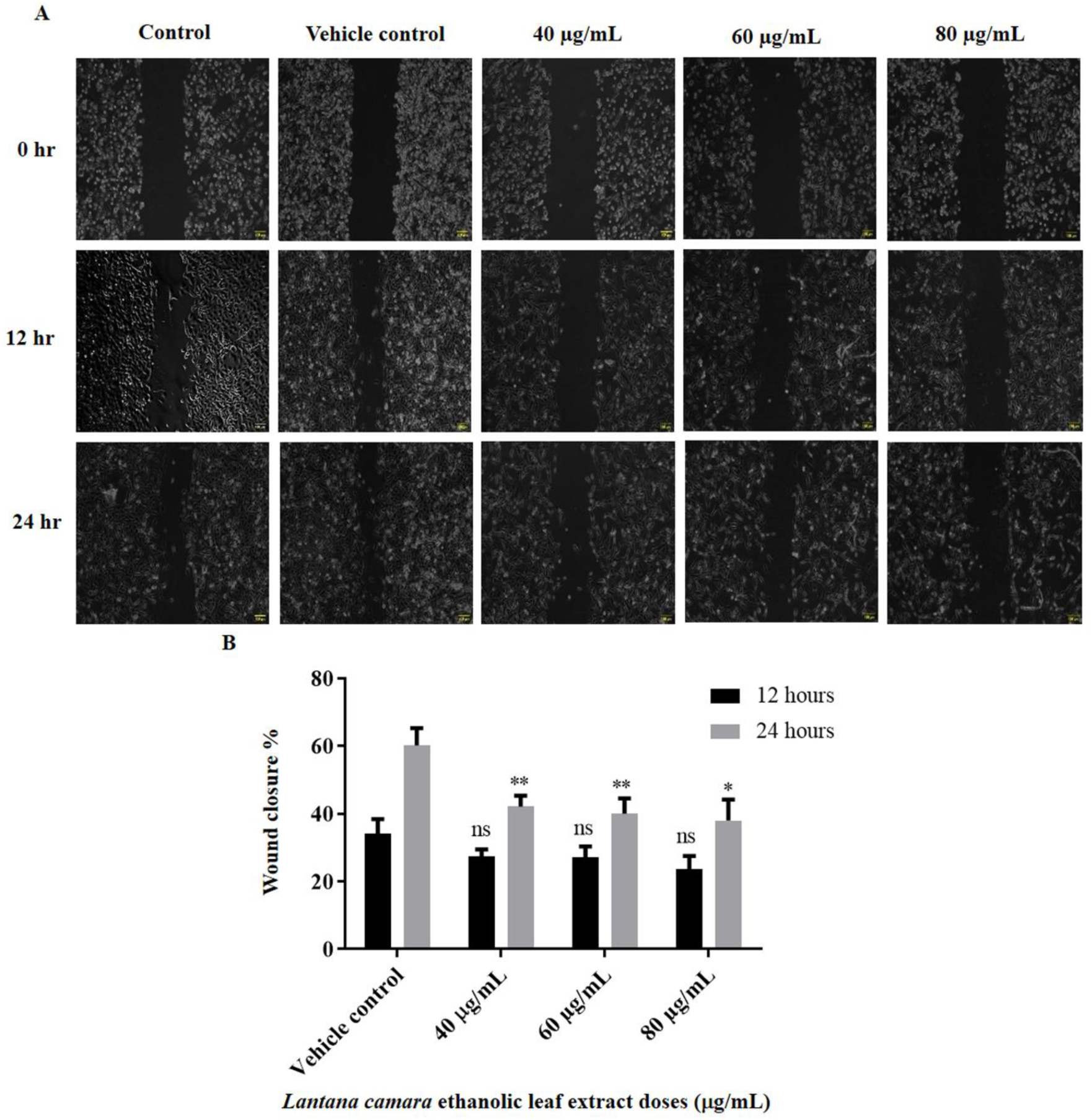
Migration of MDA-MB-231 cells was impeded by *L. camara* leaf ethanolic extract. **(A)** For wound healing assay untreated, vehicle treated and extract treated MDA-MB-231 cells were subjected to scratch at monolayer condition and then incubated for another 24 hours. Wound closure was visualized with a phase contrast microscope and images were captured at specified time intervals. **(B)** Bar diagram representing wound closure potential of MDA-MB-231 cells at different time intervals after being treated with various doses of the extract. Data is represented as means ± SEM of three independent biological replicates where * is (*P*≤0.05), ** (*P*≤0.01), *** (*P*≤0.001) and ns signifies non-significant.

### 3.6 *L. camara* leaf ethanolic extract altered the mRNA expression of key genes involved in G0/G1 cell cycle arrest, apoptosis and cell migration

RT-qPCR data obtained from *L. camara* leaf ethanolic extract-treated MDA-MB-231 cells revealed that the extract altered the mRNA levels of some key genes involved in G0/G1 cell cycle arrest, apoptosis, and cell migration.

Cyclin D1 (CCND1) is an important protein that positively regulates the progression of the cells during the G1 phase of the cell cycle (Barnes and Gillett, 1998). Whereas, p21 (CDKN1A) acts by inhibiting CDK4/6-Cyclin D complex and induces cell cycle arrest in the G1 phase (Xia et al., 2023). The expression of the Cyclin D1 (Figure 6A) gene was shown to be significantly downregulated to 0.63 ± 0.07, 0.22 ± 0.02, and 0.17 ± 0.01-folds when the MDA-MB-231 cells were treated with 40 µg/mL, 60 µg/mL and 120 µg/mL doses of *L. camara* leaf ethanolic extract, respectively compared to vehicle control. Whereas, the mRNA expression of cell cycle inhibitor p21 (Figure 6A) significantly increased 3.41 ± 0.32, 3.74 ± 0.47, and 7.55 ± 0.22-folds when treated with 40 µg/mL, 60 µg/mL and 120 µg/mL doses of the extract, respectively compared to vehicle control.

**Figure 6.**
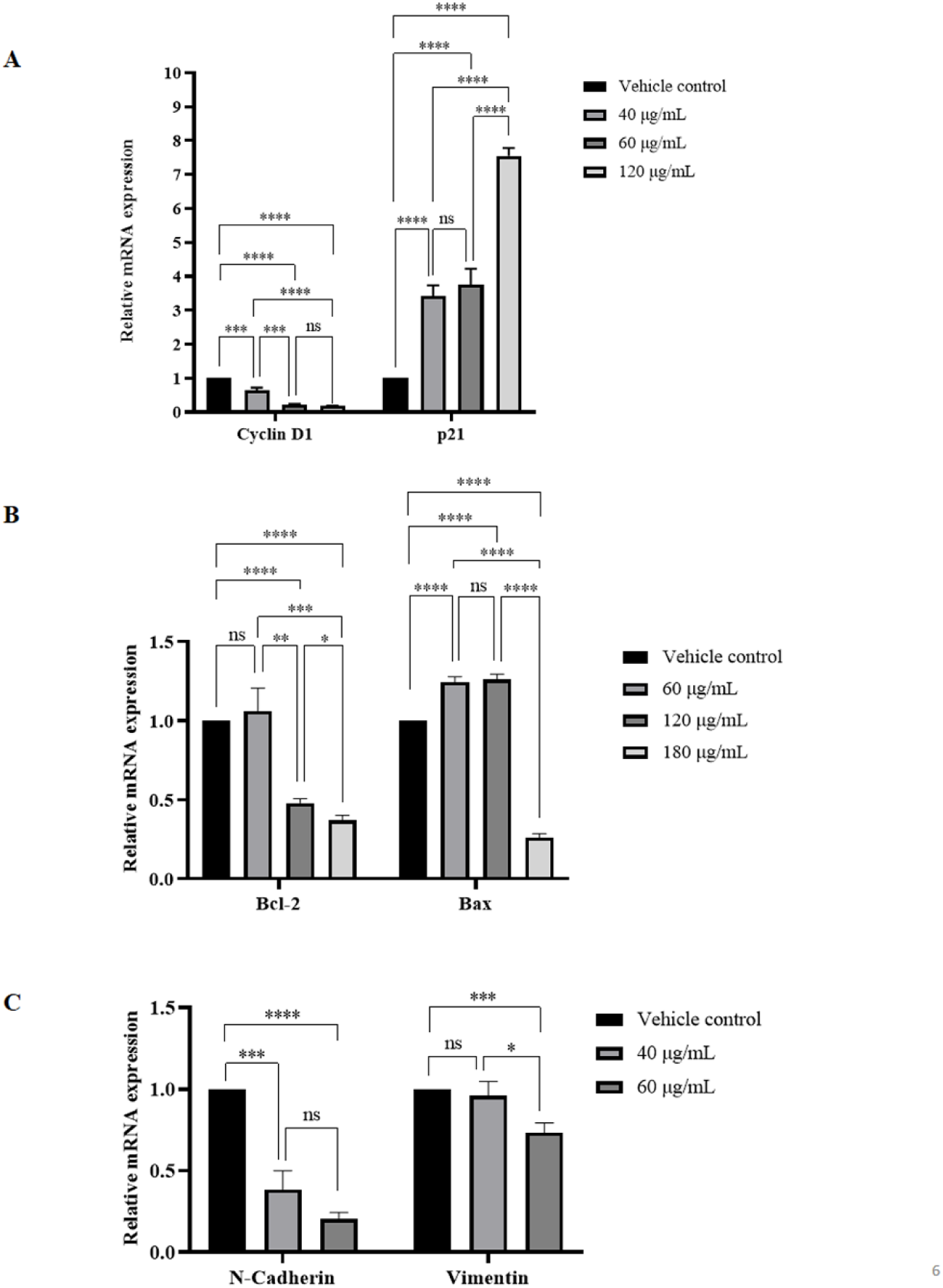
mRNA expression levels of some key genes were altered in MDA-MB-231 cells with respect to vehicle control when treated with various doses of *L. camara* leaf ethanolic extract for 24 hours. (A) Relative expression of cell cycle regulators Cyclin D1 and p21. **(B)** Relative expression of apoptosis regulators Bcl-2 and Bax. **(C)** Relative expression of cell migration regulators N-cadherin and Vimentin. Data is represented as mean ± SEM of three independent biological replicates where * is (*P*≤0.05), ** is (*P*≤0.01), *** is (*P*≤0.001), **** is (*P*≤0.0001), and ns signifies non-significant.

Bcl-2 and Bax are important members of the Bcl-2 family proteins that regulate the intrinsic apoptotic pathway (Hardwick and Soane, 2013). Bcl-2 plays an anti-apoptotic role, whereas Bax functions as a pro-apoptotic protein (Qian et al., 2022). The mRNA expression of Bcl-2 (Figure 6B) significantly decreased to 0.48 ± 0.02 and 0.36 ± 0.03-folds when treated with 120 µg/mL and 180 µg/mL doses of *L. camara* leaf ethanolic extract respectively, although no significant change was observed in case of treatment with 60 µg/mL dose of the extract. On the other hand, the transcript level of Bax (Figure 6B) increased to 1.24 ± 0.03 and 1.26 ± 0.03-folds in case of 60 µg/mL and 120 µg/mL doses of the extract, however, interestingly the mRNA level dropped to 0.26 ± 0.02-folds when treated with 180 µg/mL dose of the extract.

N-cadherin and Vimentin are mesenchymal markers that help in the epithelial-to-mesenchymal transition (EMT) of breast cancer cells (Liu et al., 2016). Upregulation of these two genes is often observed in cases of triple negative breast cancer that helps in tumor invasion and metastasis (Kvokačková et al., 2021). The mRNA expression of N-cadherin (Figure 6C) was found to be significantly downregulated to 0.38 ± 0.11 and 0.20 ± 0.04-folds when the MDA-MB-231 cells were treated with 40 µg/mL and 60 µg/mL doses of *L. camara* leaf ethanolic extract, respectively. Similarly, Figure 6C also depicts that 60 µg/mL dose of the extract significantly decreased the transcript level of Vimentin to 0.73 ± 0.06-fold, but no significant alteration was observed in case of 40 µg/mL dose of the extract. Taken together, our gene expression data positively correlates with our previous findings that *Lantana camara* ethanolic leaf extract induces cell cycle arrest at G0/G1 phase, apoptosis and inhibits cell migration of MDA-MB-231 triple-negative breast cancer cells.

## 4. Discussion

Treatment of triple negative breast cancer is challenging as it is generally impervious to conventional hormone therapy (Neophytou et al., 2018). On the other hand, the efficacy of chemotherapy is often plagued by undesirable side effects, some of which often lead to fatal consequences. Around the globe, *Lantana camara* is extensively used to treat several disorders such as fever, asthma, chicken pox, rheumatism, ulcers and many others (Chhabra et al., 1993; Taoubis et al., 1997). The bioactive constituents of *L. camara* like triterpenoids have been reported to exert cytotoxic effects at cellular levels on various cancers including breast cancer (Inada et al., 1997; Shamsee et al., 2019). However, the knowledge about how *L. camara* affects MDA-MB-231 cells (a triple negative breast cancer cell line) is largely obscure.

Here in this study, we have observed that *L. camara* leaf ethanolic extract produced morphological changes in MDA-MB-231 cells (Figure 1A). The noticeable alteration in the morphologies of the cells in the presence of the extract was indicative of cytotoxic stimulation. The experimental results from cell viability assays ascertained that the cytotoxic effect was mediated by a decline in the viability of the cells (Figure 1B). From the MTT assay result, the IC_50_ value of the leaf extract was determined which was found to be ∼111.33 μg/mL. It is to be mentioned that the extract used in this study exhibited less cytotoxic activity towards the normal kidney epithelial-like cell line HEK293T under similar experimental conditions (Supplementary Figure S2). However, when another breast cancer cell line MCF-7 was treated with the extract under similar experimental conditions, comparable cytotoxicity was observed with respect to MDA-MB-231 cells (Supplementary Figure S3). The balance between cellular proliferation and progression through the cell cycle is intricately maintained under normal physiological conditions. The inhibitory effect of the extract on the proliferation of MDA-MB-231 cells was found to be correlated with the induction of cell cycle arrest (Figure 2A and 2B) at the G0/G1 stage. The accumulation of cells in the G0/G1 phase was found to be nearly constant irrespective of the dose of the extract. This induction of cell cycle arrest was also supported by the alteration of mRNA levels of important regulators of the G0/G1 phase of the cell cycle. It was found that the mRNA level of Cyclin D1 decreased and the transcript level of p21 increased under the treatment conditions. Regulation of cell cycle and cell death have overlapping effectors; hence induction of prolonged cell cycle arrest often leads to cell death in some cases. Our study showed that the sub-G1 population was increasing slightly upon treatment with the extract. Although the observed increase in the sub-G1 population was insignificant, staining the extract treated MDA-MB-231 cells with DAPI exhibited increase in nuclear condensation at sub-IC_50_ dose (60 μg/mL) whereas fragmentation was observed at higher doses such as 120 μg/mL, 150 μg/mL and 180 μg/mL (Figure 3). Several studies have reported that chromatin condensation followed by fragmentation is a signature of various forms of cell death including apoptosis (Fatokun et al., 2014; Lacey et al., 2018; Yan et al., 2020)

The cumulative interpretation from our data exhibiting growth-inhibitory, cell cycle arrest inducing and chromatin fragmentation effects directed us to further delineate the mode of cell death. *L. camara* leaf ethanolic extract treated MDA-MB-231 cells were double stained using Annexin V-FITC/PI, and it was observed that at near and above IC_50_ doses (120 μg/mL and 180 μg/mL) the plant extract induced apoptotic as well as non-apoptotic cell death (Figure 4A and 4B). However, at sub-IC_50_ dose (60 μg/mL) there was no noticeable increase in apoptotic or non-apoptotic population when compared to the vehicle-treated cells. To further substantiate the mode of cell death, the mRNA expression levels of anti-apoptotic Bcl-2 and pro-apoptotic Bax genes were checked. It was found that the expression of Bcl-2 was downregulated in a dose-dependent manner and the expression of Bax increased in case of 60 μg/mL and 120 μg/mL doses of *L. camara* leaf ethanolic extract. Interestingly, the expression of Bax decreased under 180 μg/mL treatment condition. However, this finding is not surprising as it was reported that an increase in the Bax expression is an early event of apoptosis and in the late apoptotic phase Bax is probably degraded (Boersma et al., 1997).

Intracellular ROS generation has also been checked in MDA-MB-231 cells treated with various doses (40 μg/mL -180 μg/mL) of *L. camara* leaf ethanolic extract for 24 hours (Supplementary Figure S4) and it was observed that intracellular ROS production increased by ∼1.53 and ∼1.68 folds in MDA-MB-231 cells when treated with 120 μg/mL and 150 μg/mL doses of the extract compared to the vehicle control. This increase in ROS generation may be associated with the induction of early apoptosis under the same experimental conditions. Although this preliminary result indicates the role of ROS in the induction of apoptosis in MDA-MB-231 cells, it needs to be further validated.

The ability of cancer cells to migrate is crucial for tumor (Mehlen and Puisieux, 2006). Potential therapeutic strategies are directed towards restricting the motility of cancer cells. MDA-MB-231 cells being an aggressive metastatic TNBC cell line have an inherent capability to migrate (Huang et al., 2020). Hence, the efficacy of the extract on the migratory capacity of the cells was evaluated in this study. For this experimental purpose, cells were mainly exposed to the sub-IC_50_ doses to invalidate any prominent cytotoxic effect being induced by the extract, which in turn might alter the wound area. We could establish through our results that the leaf extract decreased the wound healing capacity of the cells, mainly after 24 hours of treatment irrespective of the doses used. To further validate the anti-migratory potential of *L. camara* leaf ethanolic extract, transcript levels of N-cadherin and vimentin (EMT markers) were checked and it was found that the mRNA expression of both the genes decreased when treated with the extract. Our finding hence established that *L. camara* leaf ethanolic extract suppressed the migratory potential of MDA-MB-231 cells.

Various chemotherapeutic drugs like Paclitaxel, Vincristine, and Sulforaphane, which have their origin in plants, have been proven effective against cancer (Greenwell and Rahman, 2015). Natural products, e.g. curcumin, genistein, resveratrol, lycopene have exhibited their cytotoxic and anti-migratory effects on triple negative breast cancer (Ke et al., 2022). The present study has elucidated that *Lantana camara* is capable of combating the aggressive nature of MDA-MB-231 cells by exhibiting a dual role in triggering cellular death and hindering the motility of the cells as well. The antiproliferative effect of natural plant products is mainly attributed to the presence of bioactive compounds such as flavonoids, terpenoids, phenols (Abotaleb et al., 2020; Kamran et al., 2022; Kopustinskiene et al., 2020). In preliminary chemical analyses, it was found that *Lantana camara* leaf ethanolic extract used in this study contains terpenoids, flavonoids, phenols and steroids (Supplementary Table S2). Although in comparison to the purified natural products like doxorubicin (Wang et al., 2020) and Paclitaxel (Vijayan et al., 2018), the cytotoxic activity of the ethanolic leaf extract used in this study showed lower activity (higher IC_50_ value), it can be speculated that the purified bioactive compound present in this leaf extract or any nano-conjugate of it will exhibit comparable cytotoxicity with respect to chemotherapeutic agents used in present days. Hence, further isolation and identification of the bioactive compounds in *L. camara* leaf ethanolic extract and studying their implications on triple negative cancer cells and *in vivo* models would provide a future platform for the area of breast cancer therapy, and will also reveal the potential and drawbacks of the leaf extract as a future therapeutic agent against breast cancer. In a recent study, it was found that a semi-synthetically modified compound of Lantadene B could reduce tumor incidence, tumor volume, and tumor burden of Swiss albino mice (Chauhan et al., 2023). In another study, it was reported that *Lantana camara* methanolic leaf extract could repress skin papilloma and reduce the death rate in Swiss albino mice (M Sharma et al., 2007). However, it has also been observed that Lantadenes can cause weight loss, hepatotoxicity, and nephrotoxicity in guineapigs (Kumar et al., 2018) and Pass et al. reported that Lantadene A could induce acute hepatotoxicity in sheep at higher doses (Pass et al., 1979). Another study revealed that the methanolic extract of *Lantana camara* leaf can cause weight loss in female mice and organ (heart and kidney) mass loss in male mice (Pour et al., 2011), but no such reports are available for humans. In humans, *Lantana camara* has only shown mild symptoms like vomiting, diarrhoea and stomach ache among children ((Carstairs et al., 2010). However, derivatization of the bioactive compounds can reduce the level of toxicity. *In silico* studies by Chauhan et al. reported that Lantadenes and their derivatives are safe and non-toxic in humans (Chauhan et al., 2023). Another drawback of plant extracts is their formulation and delivery due to low stability, poor solubility, and limited bioavailability. However, these limitations can be overcome by nano-based drug delivery systems or chemical derivatization to enhance the drug dissolution, bioavailability and formulation (Elkordy et al., 2021; Yap et al., 2021). In a recent study, Shukla et al. reported that cetuximab functionalized albumin nanoparticles of *Lantana camara* component Oleanolic acid could exert better cytotoxicity and also showed the potential for site-specific targeted delivery, bioavailability and biocompatibility in lung cancer (Shukla et al., 2023).

Taken together, the results suggest that *L. camara* leaf ethanolic extract exerts its cytotoxic effects by induction of cell cycle arrest in the G0/G1 phase and apoptosis in triple negative MDA-MB-231 cells. The extract could also impede the migration of MDA-MB-231 cells. Essential oils extracted from *Lantana camara* leaves were reported to inhibit the proliferation of different cancer cell lines by suppressing the NF-kB pathway (Sajid et al., 2021). In another study, it was reported that methanolic extract of *L. camara* leaves inhibited the proliferation of Caco cells by downregulating the PI3K/Akt signalling pathway and repressing Cyclin D1 levels via the activation of GSK-3β (El-Din et al., 2022). Molecular docking studies performed by Chauhan and coworkers found that lantadenes and their derivatives have good binding affinity towards NF-kB and IKK-beta, and this binding is responsible for the cytotoxic activity of these compounds in A375 and A431 cell lines (Chauhan et al., 2023). Therefore, it is important to delineate the signalling pathways behind the cytotoxic and anti-migratory activities of *Lantana camara* leaf ethanolic extract used in this study to understand the mode of action of the extract in MDA-MB-231 triple negative breast cancer cell line. It is known that various signalling pathways play important roles in the regulation of tumor microenvironment and natural product-mediated alterations of the tumor microenvironment is a key aspect of cancer immunotherapy (Zhang et al., 2022). Thus the leaf extract used in this study or specific bioactive compounds may also have the property to modulate the tumor microenvironment. It was reported that different natural compounds can act synergistically with several existing chemotherapeutic agents to enhance their efficacy (Castañeda et al., 2022). It was also reported that natural products like polyphenols, alkaloids, terpenoids, saponins, and bioactive polysaccharides have the potential to be used as adjuvant therapy and cancer immunotherapy (Dong et al., 2022). Other studies have also found that *Lantana camara* methanolic leaf extract and Lantadene A and its congeners possess chemopreventive activity against skin cancer (Manu Sharma et al., 2007; M Sharma et al., 2007). A comparison of activities of different *Lantana camara* extracts, bioactive compounds and related nanoparticles is presented in Table S3 in the Supplementary section. A detailed understanding of the signalling pathway and isolation of the bioactive compounds present in the extract will pave the path for studying the synergistic effect of the extract with different conventional chemotherapeutic agents, and will also reveal the potential of the extract to be used as adjuvant therapy and as a chemopreventive agent for triple negative breast cancer.

## Conclusion

*Lantana camara* leaf ethanolic extract is cytotoxic to the triple negative breast cancer cell line, MDA-MB-231. The extract induces G0/G1 cell cycle arrest and DNA condensation ultimately leading to apoptosis-mediated cellular death. In conclusion, the growth inhibitory effect of the extract could be largely attributed to its ability to induce cell cycle arrest and at the same time its potential to initiate apoptosis. Furthermore, the migratory potential of the cells was also reduced in the presence of the extract. Thus, our findings are suggestive of the fact that *Lantana camara* is a potential source of chemotherapeutic agents for combating triple negative breast cancer. Future works focusing on the molecular mechanisms and signalling pathways behind cellular death and decreased migration would further enrich the understanding of the mode of action of this plant extract. However, identification of the bioactive compounds present in the extract is essential which would enable researchers to expand the repertoire of potential source of chemotherapeutic agents.

## Supporting information

Supplementary Section

## Acknowledgements

All authors would like to acknowledge Prof. Subhajit Bandyopadhyay for providing rotary-evaporator facility. The authors would also like to acknowledge Mr. Ritabrata Ghosh and Mr. Tamal Ghosh for their technical support in epifluorescence microscopy and flow-cytometry respectively. All authors would also like to convey their gratitude to Professor Mushfiquddin Khan, Department of Paediatrics, Medical University of South Carolina. AP and SS acknowledge the Department of Biotechnology, Government of India (DBT-India) for research fellowship. SD acknowledges IISER Kolkata for the Postdoctoral Fellowship.

## Author Contributions

AP, SS, SD and TKS conceived the study and designed the experiments. AP, SS and SD did the experiments. AP, SS, SD and TKS analyzed the data and wrote the manuscript.

## Availability of Data and Materials

In this study, all data generated or analyzed are included in the article.

## Ethical Approval

Not applicable.

## Consent to Participate

All authors had consented to participate in the study.

## Consent for Publication

All authors have given consent for publication.

