## Supplementary Section for "Effect of *Lantana camara* ethanolic leaf extract on survival and migration of MDA-MB-231 triple negative breast cancer cell line"

**Solvent (Ethanol) tolerance assay:**

To assess the effect of the solvent (Ethanol) on the MDA-MB-231 cell line, a solvent tolerance study was performed. *Lantana camara* leaves were extracted using absolute molecular grade ethanol (Merck, Germany) and subsequent working stocks of the *Lantana camara* extract were also prepared in the molecular grade ethanol. For this reason, a solvent tolerance assay was necessary to select a particular concentration (percentage) of the solvent as vehicle that will cause a minimum effect on the cells. MDA-MB-231 cells were seeded in 96 well plate at a density of 0.5x10^4^ cells/well and were incubated overnight at 37ºC in 95% air in a humidified incubator with 5% CO_2_. Cells were then treated with different percentages of ethanol (0.2% to 6%) and untreated MDA-MB-231 cells served as control. Cells were then incubated for 24 hours. After 24 hours, the cytotoxic effect of ethanol was determined using MTT colourimetric assay following the same protocol as mentioned in the material and method section (2.4) of the manuscript.

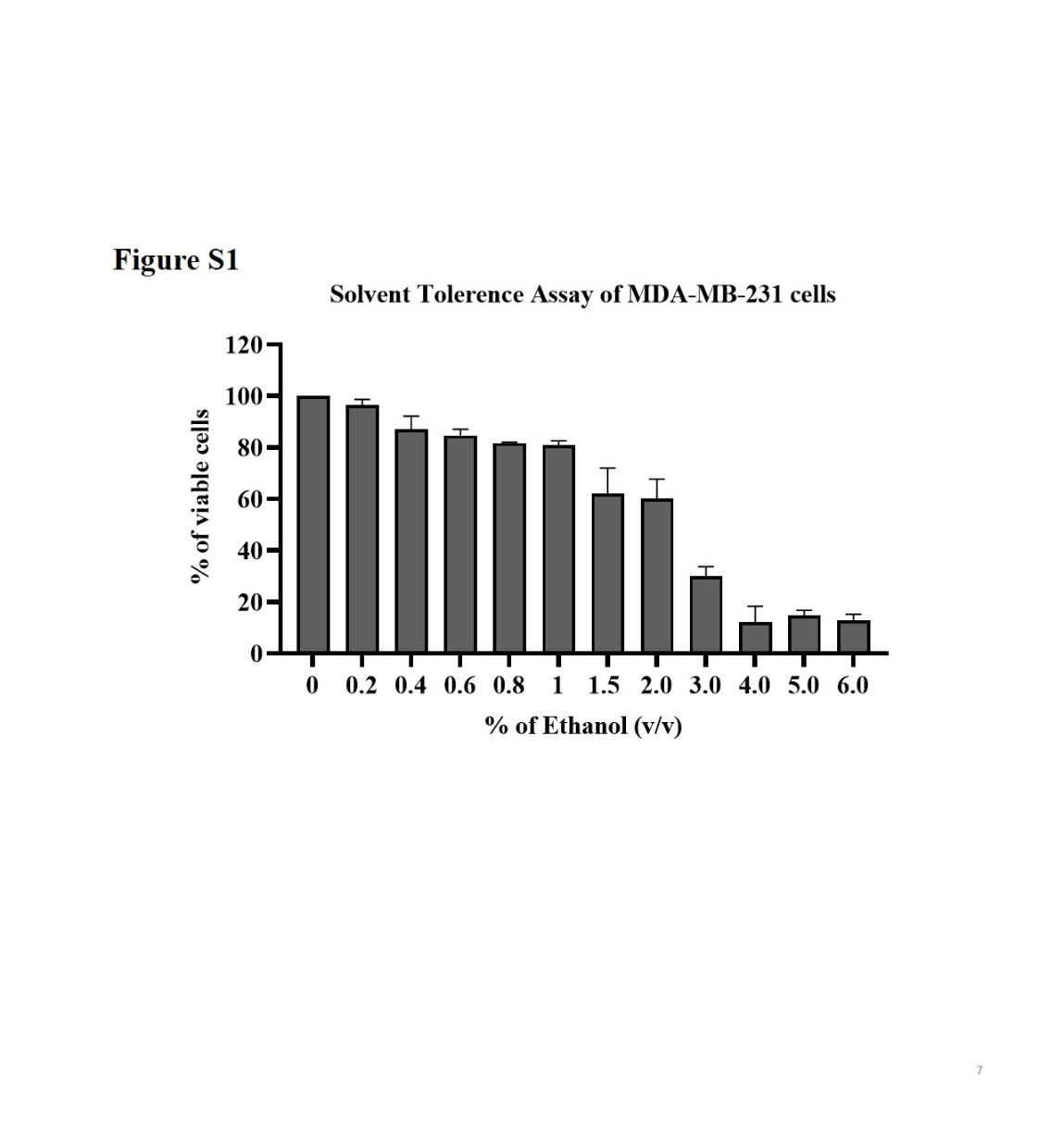

**Figure S1:** Solvent tolerance assay of MDA-MB-231 cells treated with various concentrations (% v/v) of ethanol for 24 hours. Data is represented as Mean ± SEM of three independent biological replicates.

**Result**: There was a noticeable decrease in the viability of MDA-MB-231 cells with the increase in solvent percentage at higher concentrations (1.5% - 6%). However, at 0.2% - 1% solvent concentration, the decrease in cell viability was minimal. From the above range, 0.8% ethanol was selected as vehicle control due to its minimal cytotoxic effect on MD-MB-231 cells and as an appropriate concentration for leaf extract solubility.

**Table S1.** List of primers used for RT-qPCR study

| **Target gene** | **Primer sequences** |
| --- | --- |
| 18S | Forward Primer: 5’-ACGGAAGGGCACCACCAGGAGT-3’  Reverse Primer: 5’-GAACGGCCATGCACCACCACC-3’ |
| Cyclin D1 (CCND1) | Forward Primer: 5’-CCCGAGGAGCTGCTGCAAAT-3’  Reverse Primer: 5’-CGAAGGTCTGCGGCTGTTTG-3’ |
| p21 (CDKN1A) | Forward Primer: 5’-CACTCAGAGGAGGCGCCATGTC-3’  Reverse Primer: 5’-ATCGCTCACGGGCCTCCTGGAT -3’ |
| Bcl-2 | Forward Primer: 5’-TCTTCAGGGACGGGGTGAACT-3’  Reverse Primer: 5’-TTCCACAAAGGCATCCCAGCC-3’ |
| Bax | Forward Primer: 5’-ACAGGGGCCCTTTTGCTTCA-3’  Reverse Primer: 5’-ACGGCGGCAATCATCCTCTG-3’ |
| N-cadherin | Forward Primer: 5’-CAGCAGCCTGACACTGTGGA-3’  Reverse Primer: 5’-GCCGCTTTAAGGCCCTCATT-3’ |
| Vimentin | Forward Primer: 5’-CAATGCGTCTCTGGCACGTC-3’  Reverse Primer: 5’-TGCAGCTCCTGGATTTCCTCT-3’ |

**Comparison of the effect of *Lantana camara* leaf ethanolic extract on non-cancer kidney epithelial-like HEK293T cells and MDA-MB-231 cells**

**Cell morphology analysis**

Morphology of the HEK293T cells was observed following 24 hours of treatment with various doses of *Lantana camara* leaf extract (40 µg/mL – 150 µg/mL) using the same protocol as mentioned in the material and method section (2.3) of the manuscript

**Trypan Blue dye exclusion assay for determining cell viability**

Trypan Blue dye exclusion assay is used to determine the number of viable cells present in a cell suspension. It is based on the principle that live cells possess intact cell membranes that exclude Trypan Blue and appear transparent under a microscope whereas dead cells do not exclude the dye and appear blue (Strober, 2015). Briefly, both HEK293T and MDA-MB-231 cells were seeded in 35mm dishes at a density of 0.15 x10^6^ cells/dish and were incubated overnight at 37ºC in 95% air in a humidified incubator with 5% CO_2_. Cells were then treated with various doses of *Lantana camara* leaf extract (40 µg/mL – 180 µg/mL) along with vehicle (0.8% ethanol) treated cells as control, and were incubated for 24 hours. Cells were then harvested using 1X trypsin-EDTA solution (HiMedia, India) and were mixed with Trypan Blue dye (HiMedia, India) in a 1:1 ratio and 10μL of cell suspension with Trypan Blue dye was transferred to each chamber of hemocytometer and were counted under microscope. The viability of the cells was determined using the following formula:

Total number of viable cells per mL of the aliquot

% Cell viability =

X 100

Total number of cells per mL of aliquot

The experiment was done in three replicates and the data was represented as mean ± SD and plotted using GraphPad Prism 6.01 (GraphPad Software, CA, USA).

**3-(4,5-dimethylthiazol-2-yl)-2,5-diphenyltetrazolium bromide (MTT) colourimetric assay for cell viability:**

MTT Assay was also performed for the above-mentioned experimental conditions using the same protocol as mentioned in the material and method section (2.4) of the manuscript.

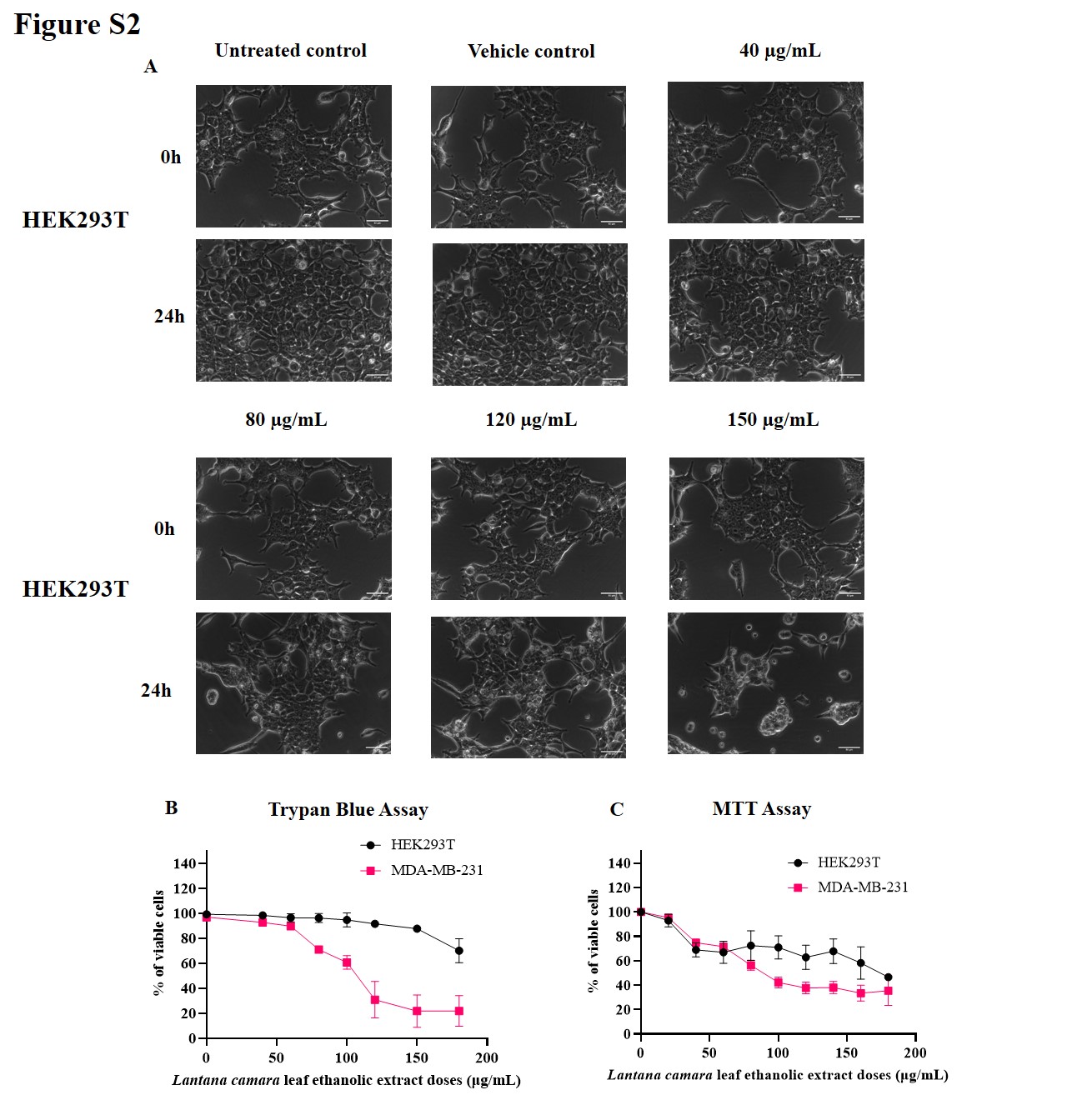

**Figure S2:** Morphology analysis of HEK293T cells (A) under phase contrast microscope (40X) after treatment with *Lantana camara* leaf extract. Trypan Blue Assay (B) of HEK293T and MDA-MB-231cells. MTT Assay (C) of HEK293T and MDA-MB-231cells. Data is represented as Mean ± SD of three independent replicates for Trypan Blue Assay and Mean ± SEM of three independent biological replicates for MTT Assay.

**Result:** Comparing the morphology analysis (A), Trypan Blue Assay (B) and MTT Assay (C) results, it was found that *Lantana camara* leaf extract is more effective towards triple negative MDA-MB-231 breast cancer cells than HEK293T cells. IC_50_ of *Lantana camara* extract for HEK293T was calculated from MTT Assay data and was found to be 182.07 μg/mL which is much higher than the IC_50_ value of the extract for MDA-MB-231 cells (IC_50_ = 111.33 μg/mL).

**Biochemical tests to identify active compounds present in *Lantana camara* leaf ethanolic extract**

**Test for Terpenoids:** Salkowski test was used to detect terpenoids. Extract (5 mL) was mixed with chloroform (2 mL), and concentrated sulphuric acid (3 mL) was then carefully added to form a layer. A reddish-brown colouration of the interface was formed to show positive results for the presence of terpenoids (Das et al., 2014).

**Test for Flavonoids:** A few drops of 1% NH3 solution were added to the plant leaf extract (0.5 g) in a tube. Yellow colouration implies that flavonoid compounds are present (Swamy et al., 2012).

**Test for Phenols:** 1 mL extract was mixed with 1 mL of distilled water. Then a few drops of neutral 5% ferric chloride solution were added. A dark green colour indicates the presence of phenolic compounds (Bargah, 2015).

**Test for Steroids:** 5 mL plant extract was taken in a test tube and mixed with chloroform (10 mL). Then equal volume of concentrated sulphuric acid was added to the test tube by the sides. If the upper layer in the test tube turns red and sulphuric acid layer shows yellow with green fluorescence, it shows the presence of steroids (Hossain et al., 2013).

**Test for Tannins:** 2 mL of the extract was stirred with 2 mL of distilled water and a few drops of ferric chloride (FeCl_3_) solution were added. The formation of a green precipitate indicates the presence of tannins (Bargah, 2015).

**Test for Saponins:** 5 mL of extract was shaken vigorously with 5 mL of distilled water in a test tube and warmed. The formation of stable foam indicates the presence of saponins (Bargah, 2015).

**Result**

**Table S2.** Phytochemical Analyses

| **Phytoconstituents** | **Presence/Absence** |
| --- | --- |
| Terpenoids | Present |
| Flavonoids | Present |
| Phenols | Present |
| Steroids | Present |
| Tannins | Absent |
| Saponins | Absent |

**Comparison of the effect of *Lantana camara* leaf ethanolic extract on MCF-7 cells and MDA-MB-231 cells**

**Trypan Blue dye exclusion assay for determining cell viability**

Trypan Blue dye exclusion assay is used to determine the number of viable cells present in a cell suspension. It is based on the principle that live cells possess intact cell membranes that exclude Trypan Blue and appear transparent under a microscope whereas dead cells do not exclude the dye and appear blue (Strober, 2015). Briefly, both MCF-7 and MDA-MB-231 cells were seeded in 35mm dishes at a density of 0.15 x10^6^ cells/dish and were incubated overnight at 37ºC in 95% air in a humidified incubator with 5% CO_2_. Cells were then treated with various doses of *Lantana camara* leaf extract (40 µg/mL – 180 µg/mL) along with vehicle (0.8% ethanol) treated cells as control, and were incubated for 24 hours. Cells were then harvested using 1X trypsin-EDTA solution (HiMedia, India) and were mixed with Trypan Blue dye (HiMedia, India) in a 1:1 ratio and 10μL of cell suspension with Trypan Blue dye was transferred to each chamber of hemocytometer and were counted under microscope. The viability of the cells was determined using the following formula:

Total number of viable cells per mL of the aliquot

% Cell viability =

X 100

Total number of cells per mL of aliquot

The experiment was done in three replicates and the data was represented as mean ± SD and plotted using GraphPad Prism 6.01 (GraphPad Software, CA, USA).

**3-(4,5-dimethylthiazol-2-yl)-2,5-diphenyltetrazolium bromide (MTT) colourimetric assay for cell viability:**

MTT Assay was also performed for the above-mentioned experimental conditions using the same protocol as mentioned in the material and method section (2.4) of the manuscript.

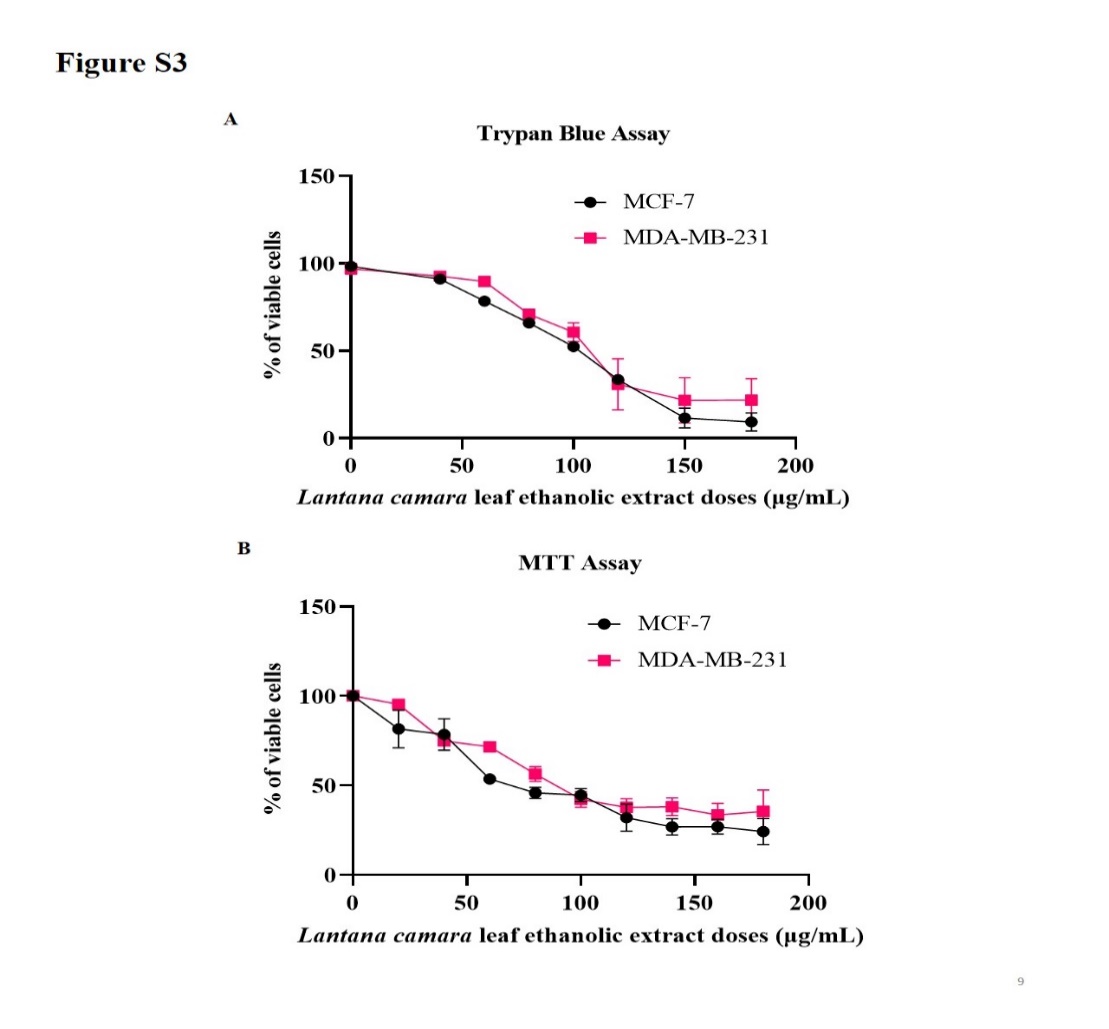

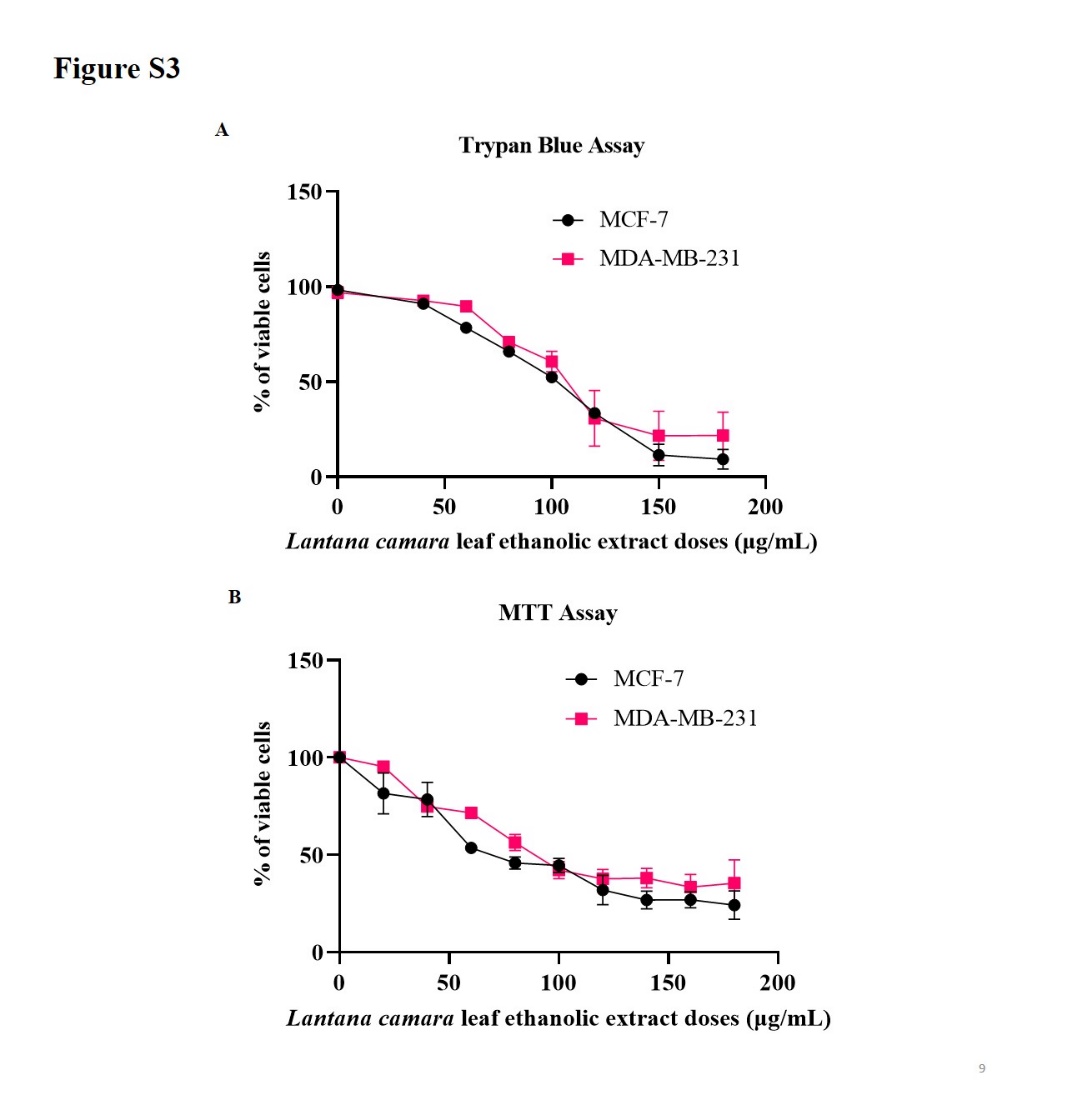

**Figure S3:** Trypan Blue Assay (A) of MCF-7 and MDA-MB-231 cells. MTT Assay (B) of MCF-7 and MDA-MB-231 cells. Data is represented as Mean ± SD of three independent replicates for Trypan Blue Assay and Mean ± SEM of three independent biological replicates for MTT Assay.

**Result:** Analyzing Trypan Blue Assay (A) and MTT Assay (B) data of MCF-7 and MDA-MB-231 cells, it has been observed that *Lantana camara* leaf ethanolic extract is almost equally cytotoxic to both the cell lines. IC_50_ of the extract was calculated for MCF-7 cells from the MTT Assay data and was found to be 93.25 µg/mL. This IC_50_ value for MCF-7 cells is comparable to the IC_50_ value of the extract for MDA-MB-231 cells (IC_50_ = 111.33 µg/mL).

**Measurement of intracellular Reactive Oxygen Species (ROS) generation**

To evaluate the role of oxidative stress behind the cytotoxic effects of *Lantana camara* leaf ethanolic extract, intracellular ROS production was measured by using 2,7 Dichlorofluorescein Diacetate (DCFH, SRL, India). DCFH is a cell-permeable dye that diffuses into the cells, gets deacetylated by cellular esterases into a non-fluorescent compound, and then gets oxidized by ROS into highly fluorescent 2,7-Dichlorofluorescein (DCF) (Marvibaigi et al., 2016). Briefly, MDA-MB-231 cells were seeded in a 24-well plate with a density of 1x10^5^ cells/well and incubated overnight at 37°C in 5% CO_2_. After incubation, cells were treated with different concentrations of *Lantana camara* leaf ethanolic extract (40 µg/mL-180 µg/mL), along with vehicle control (0.8% ethanol) and 500 µM H_2_O_2_ (as positive control) for 24 hours. After 24 hours, cells were harvested, washed with 1X PBS, and re-suspended in 10 µM DCFH solution (20 mM DCFH stock solution was diluted in pre-warmed serum-free DMEM to prepare 10 µM DCFH working solution). Cells were then incubated at 37°C for 30 minutes. Then the cells were washed twice with 1X PBS. Finally, the cells were resuspended in 1X PBS and analysed using BD FACS Fortessa flow cytometer (BD Biosciences, USA) at 488 nm and 530 nm excitation and emission, respectively. The fluorescence intensity of DCF is directly proportional to intracellular ROS generation and the graph was plotted as fold change relative to the vehicle control. The experiment was done in three replicates and the data was represented as mean ± SD and plotted using GraphPad Prism 6.01 (GraphPad Software, CA, USA).

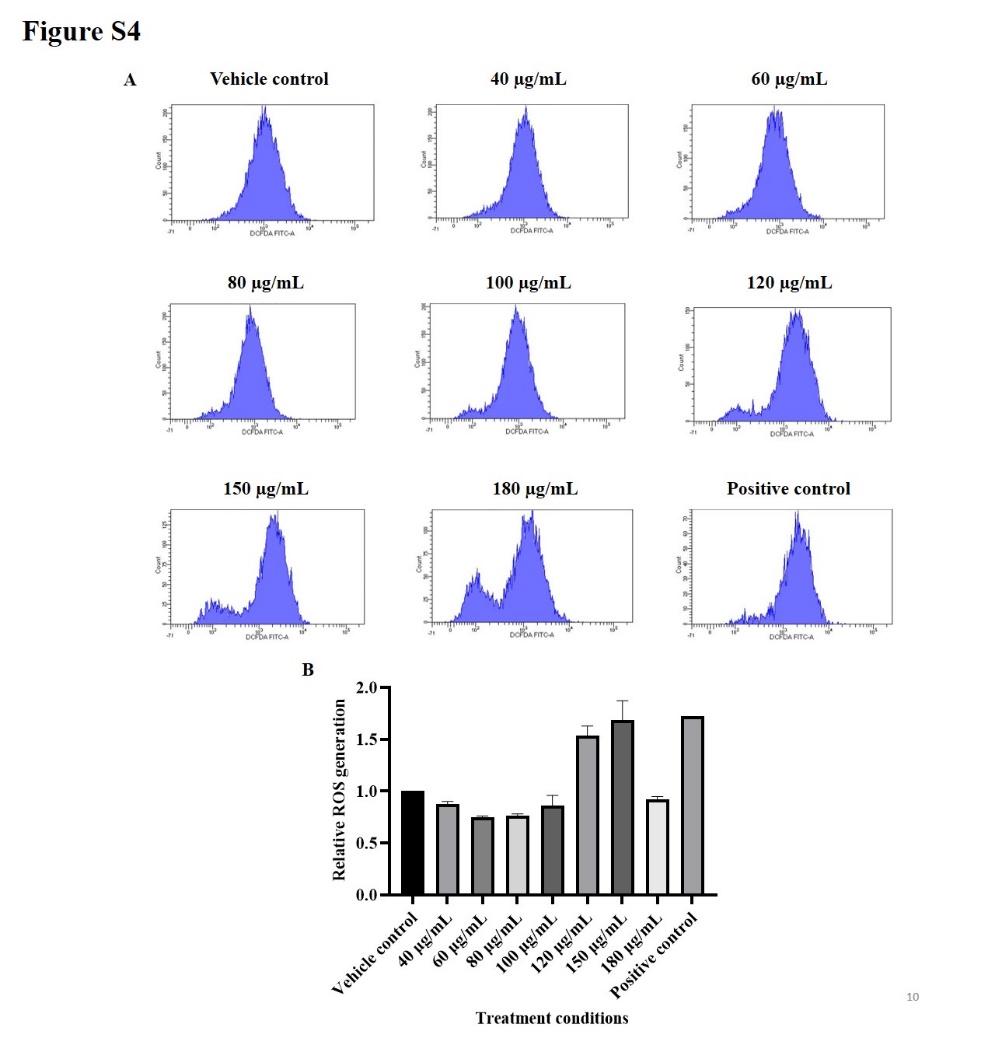

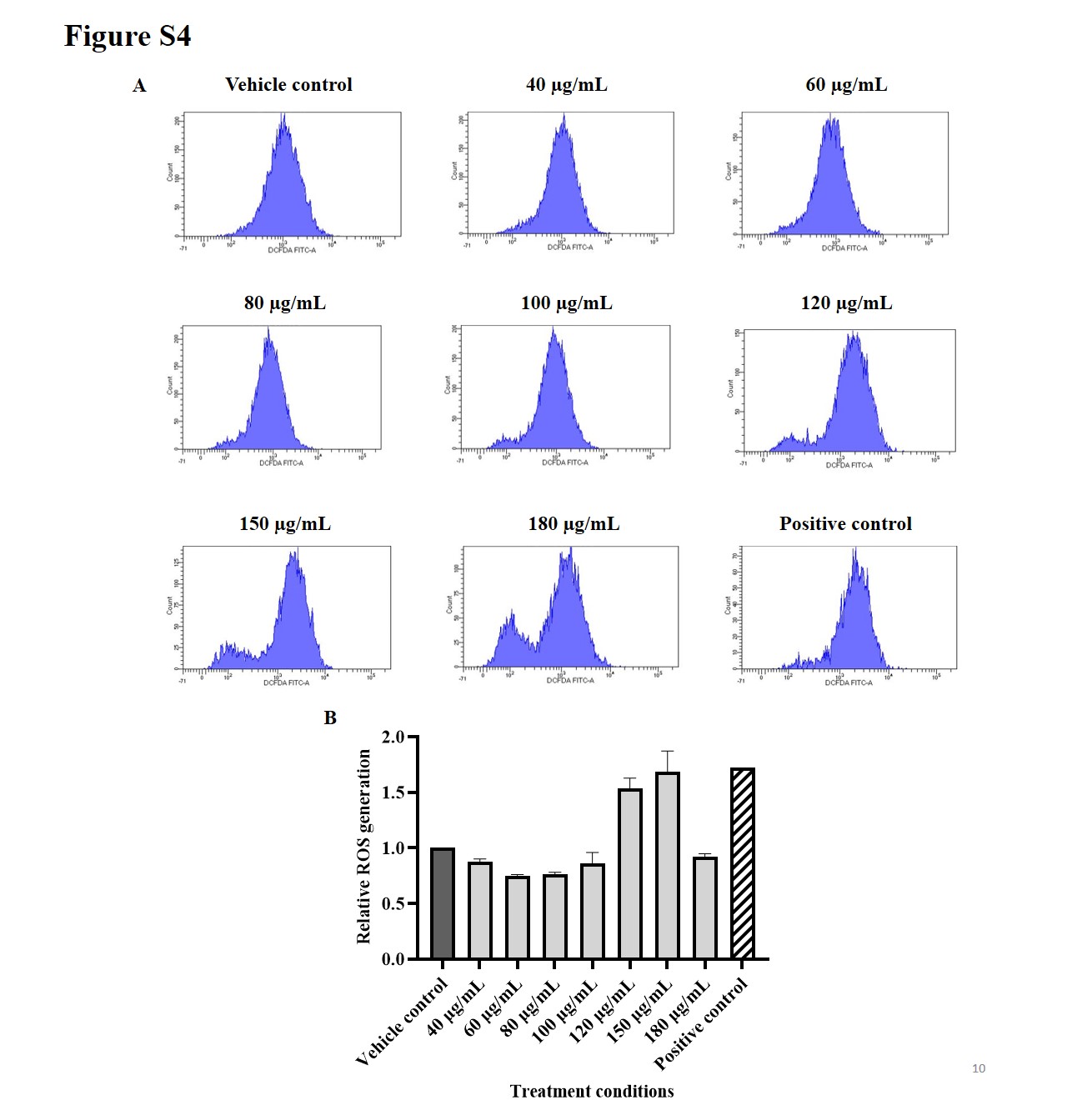

**Figure S4:** (A) Representative images of measurement of intracellular Reactive Oxygen Species (ROS) generation upon 24 hours of treatment with *Lantana camara* extract (40 µg/mL – 180 µg/mL) and 500 µM H_2_O_2_ (as positive control). (B) Bar diagram showing relative ROS generation to vehicle control under different treatment conditions. Data is represented as Mean ± SD of three independent replicates.

**Result:** From the above data, it was observed that there was no noticeable difference in the production of intracellular ROS from 40 μg/mL to 100 μg/mL dose of the extract. However, intracellular ROS increased by 1.53 and 1.68 folds in MDA-MB-231 cells when treated with 120 μg/mL and 150 μg/mL of leaf extract respectively for 24 hours as compared to the control condition (vehicle control). These increases in ROS generation indicate that it could be associated with the induction of early apoptosis under the same experimental conditions.

**Table S3.** Cytotoxic activity of *Lantana camara* extract/bioactive compounds against different cancer cell lines

| **Type of *Lantana camara* extract/Bioactive compounds isolated from *Lantana camara*** | **Cancer cell line** | **Outcome** | **IC_50_ Value** | **References** |
| --- | --- | --- | --- | --- |
| Methanolic extract from different parts of *Lantana camara* | A549, B16-F10, HEp-2 | Leaf extract exhibited comparatively more cytotoxic activity against all the cell lines | - A549: 58.48±2.83 µg/mL - B16-F10: 243.38±8.46 µg/m - HEp-2: 221.18±3.94 µg/mL | (Hariharapura et al., 2004) |
| *Lantana camara* ethanol extract (95%) | MCF-7 | - Cytotoxic - Induced apoptosis - Downregulation of Bcl-2 and upregulation of Bax and Bid - Modulated cleavage of Caspase-8, Caspase-9 and PARP | In the range from 32.39µg/mL to 46.94 µg/mL | (Han et al., 2015) |
| *Lantana camara* root extract-mediated gold nanoparticles | MDA-MB-231 | - Showed *in vitro* antioxidant activity - Showed cytotoxic activity in MDA-MB-231 cells | 17.72 µg/mL | (Ramkumar et al., 2017) |

| **Type of *Lantana camara* extract/Bioactive compounds isolated from *Lantana camara*** | **Cancer cell line** | **Outcome** | **IC_50_ Value** | **References** |
| --- | --- | --- | --- | --- |
| Pentacyclic triterpenoids isolated from *Lantana camara* leaf methanolic extract | MCF-7 | - Separated pentacyclic triterpenoids (Lantadene A, B, C and icterogenin) - All the compounds showed free radical scavenging activity; Lantadene A and B are the most potent - All the compounds showed cytotoxic activity; Lantadene B is the most potent - Lantadene B induced G1 cell cycle arrest and increased Caspase-9 activity | - Lantadene A: 227 µg/mL - Lantadene B: 112.2 µg/mL - Lantadene C: 233.1 µg/mL - Icterogenin: 180.6 µg/mL | (Shamsee et al., 2019) |
| Gold nanoparticles of Lantadene A (isolated from *Lantana camara* leaf methanolic extract) | MCF-7 | - Genotoxic and cytotoxic - Downregulation of Bcl-2 and upregulation of p53 and Bax | - Lantadene A: 158.20 µg/mL - Lantadene A-loaded gold nanoparticles: 64.39 µg/mL | (Jaafar et al., 2020) |

| **Type of *Lantana camara* extract/Bioactive compounds isolated from *Lantana camara*** | **Cancer cell line** | **Outcome** | **IC_50_ Value** | **References** |
| --- | --- | --- | --- | --- |
| *Lantana camara* methanolic leaf extract | MCF-7, MDA-MB-231, Caco, PCL | - Characterized the bioactive compounds of the extract - Extract showed free radical scavenging and anti-inflammatory activities - Cytotoxic to all the cell lines - Induced cell cycle arrest in Sub-G0/G1 phase in Caco cells - Upregulation of p53, GSK-3β and downregulation of PI3K, p-Akt and Cyclin D1 in Caco cells | - MCF-7: 78.08±1.39 µg/mL - MDA-MB-231: 74.3±1.19 µg/mL - Caco: 45.65±1.64 µg/mL - PCL: 52.55±1.14 µg/mL | (El-Din et al., 2022) |
| Silver nanoparticles synthesized using *Lantana camara* leaf extract | MCF-7, A549 | - Cytotoxic to both the cell lines | - MCF-7: 46.67 µg/mL - A549: 49.52 µg/mL | (Hublikar et al., 2023) |

| **Type of *Lantana camara* extract/Bioactive compounds isolated from *Lantana camara*** | **Cancer cell line** | **Outcome** | **IC_50_ Value** | **References** |
| --- | --- | --- | --- | --- |
| Cetuximab conjugated albumin nanoparticles of oleanolic acid (isolated from *Lantana camara* root extract) | A549 | - Cytotoxic - Induced G0/G1 cell cycle arrest - Increased intracellular ROS production - Induced apoptosis | - Oleanolic acid: 98.21 ± 1.45 μg/mL - Albumin nanoparticles of oleanolic acid:13.87 ± 1.28 μg/mL - Cetuximab conjugated albumin nanoparticles of oleanolic acid: 4.34 ± 1.90 μg/mL | (Shukla et al., 2023) |
| Lantadene A and B, Reduced Lantadenes and Lantadene ester derivatives (Lantadenes isolated from *Lantana camara* leaf ethyl acetate extract and then semi-synthetically modified into other compounds) | A375, A431 | - Cytotoxic - Compounds are safe and non-toxic as determined by *in silico* studies - Compounds have good binding affinity towards NF-kB and IKK-beta as determined by molecular docking studies | - A375: ranging from 3.027 μM to 12.63 μM - A431: ranging from 12.09 μM to 45.68 μM | (Chauhan et al., 2023) |
| *Lantana camara* leaf ethanolic extract | MDA-MB-231 | - Cytotoxic activity - Induction of G0/G1 cell cycle arrest and apoptosis - Impeded cell migration - Alteration of mRNA expression of some key genes | 111.33 µg/mL | Present study |

^*^Treatment time varied from 24 hours to 72 hours for different studies.
